# Same principle, but different computations in representing time and space

**DOI:** 10.1101/2023.11.05.565686

**Authors:** Sepehr Sima, Mehdi Sanayei

## Abstract

Time and space are two intertwined contexts that frame our cognition of the world and have shared mechanisms. A well-known theory on this case is ‘A Theory of Magnitude (ATOM)’ which states that the perception of these two domains shares common mechanisms. However, evidence regarding shared computations of time and space is intermixed. To investigate this issue, we asked human subjects to reproduce time and distance intervals with saccadic eye movements in similarly designed tasks. We applied an observer model to both modalities and found underlying differences the processing of time and space. While time and space computations are both probabilistic, adding prior to space perception minimally improved model performance, as opposed to time perception which was consistently better explained by Bayesian computations. We also showed that while both measurement and motor variability were smaller in distance than time reproduction, only the motor variability was correlated between them, as both tasks used saccadic eye movements for response. Our results suggest that time and space perception abide by the same algorithm but have different computational properties.

## Introduction

The study of time perception has demonstrated the complex interplay of spatiotemporal information in the brain. As temporal processes are linked to their embodied experience of the environment ^1^, analyzing the spatial dimension in the study of time perception can provide a powerful tool for understanding the underlying mechanisms that allow us to comprehend the external world. Time and space are two aspects of the physical world that frame our experience of the world. The perception of time and space occurs in an interrelated fashion in human cognition, and recent research shows a growing interest in understanding the underlying mechanisms of their relationship ^2–5^. It has been suggested that time perception occurs through spatialization of time intervals in the face of movement-based events of the world ^6^. Such spatialization could be seen in the way we have tied the perception of time to space by devising various types of clocks to keep track of the passage of time.

In modern science, time and space have been represented with measurable proxies imbued with operational definitions, i.e., time and distance intervals, respectively ^7^, which provide a framework for studying and quantifying these abstract concepts. This has made possible the study of the relationship between time and space perception. A variety of time-space interactions in the human perceptual system has been observed. For example, spatiotemporal interference is one such interaction where spatial information can distort the perception of temporal information and vice versa ^8^. Peri-saccadic spatiotemporal compression is another phenomenon that serves as evidence of common mechanisms in the perception of time and space ^9^.

A Theory of Magnitude (ATOM) proposed by Walsh ^10^ suggests that the brain has a core common magnitude system for time, space, and quantity. According to this theory, the neural mechanisms underlying the perception of time, space, and quantity are intertwined, and the brain processes these dimensions in a unified manner. This theory has been supported by empirical evidence from studies that have shown that the perception of time and space share common neural substrates ^11–16^. A recent meta-analysis of neuroimaging studies ^17^ has suggested that there is a common system of brain regions that are activated during both time and space processing, including bilateral insula, the pre-supplementary motor area (pre-SMA), the right frontal operculum, and intraparietal sulci. At the neuronal level, it has been observed that spatial information could at least be partially derived from temporal information ^18^.

Despite these findings, the precise nature and the extent to which these perceptual domains share common mechanisms remain unknown. This has spurred several investigations into better understanding the relationship between time and space ^19,20^. Most studies to date have approached the question in terms of the interferences that occur between perceptual domains ^21–23^. We attempted to approach this question by utilizing behavioral modeling and model comparison.

A Bayesian understanding of timing has revealed that the interaction of the temporal context and the internal ongoing processes culminates in the calibration of estimated intervals ^24^ in the form of perceptual biases. Such a formulation of interval timing presents us with two stages in the process of timing, i.e., the measurement (perception) and the reproduction (action) phases of interval timing ^25^. The link between the measurement and the reproduction is actualized by an estimation function in the observer model. Based on the nature of the observer model (ideal vs non-ideal), the estimation functions differ. Bayesian least squares (BLS) and maximum likelihood estimation (MLE) estimators have been used in the literature as prior-dependent and prior-independent functions, respectively ^25^. The Bayesian perspective has also been explored in human spatial navigation ^26,27^. Thus, the probabilistic nature of spatiotemporal information could be well captured by an optimum-seeking system which combines experience-dependent information with contextual noisy measurements to generate an estimate of various facets of time and space. It remains unclear whether the perceptual biases in time and space are both attributable to the prior information.

In this study, we compared the how spatial and temporal measurements are implemented by probing sources of variability in the process of time and distance measurement and reproduction, within a probabilistic framework. We showed that the perceptual biases in time perception are explained by prior-dependent computations, as previously shown in the literature. On the other hand, we cast doubt on the contribution of prior information to the observed perceptual biases in space perception.

## Results

### Bayesian Observer modeling of time and distance reproduction tasks

We plotted subjects’ reproduced time and distance as a function of the presented duration and distance (Fig. 2). In both time and distance, we observed that reproduced time and distance roughly followed the presented time and distance, respectively. In order to observe the goodness-of-fit across our four models (BLS_2p_, BLS_3p_, MLE_2p_, MLE_3p_), we fitted these models to our data, separately, and plotted simulations from the best fitted models on our data. As it is visually evident, the BLS models outperformed MLE models in the time domain. In the distance domain, MLE and BLS models were very similar. We calculated Akaike information criterion (AIC), Bayesian information criterion (BIC), and cross-validated log likelihood (CLL) across subjects for each model. We considered a value of 5 in ‘2 × difference in CLL’ as the cutoff point in model comparison for each pair as was previously suggested. In the time domain, we found that the BLS_3p_ model is a better fit for 11/20 subjects compared to the other three models (for CLL values, and summary of result, see Table 1). To further investigate the validity of such results, we ran pairwise comparisons of models’ CLL values for each subject. These analyses revealed that BLS_2p_ is a better fit than MLE_2p_ in 17/20 subjects (Fig. 3A), BLS_3p_ is a better fit than MLE_3p_ in 19/20 subjects (Fig. 3B), and BLS_3p_ is a better fit than BLS_2p_ in 10/20 subjects (Fig. 3C).

**Figure 1.**
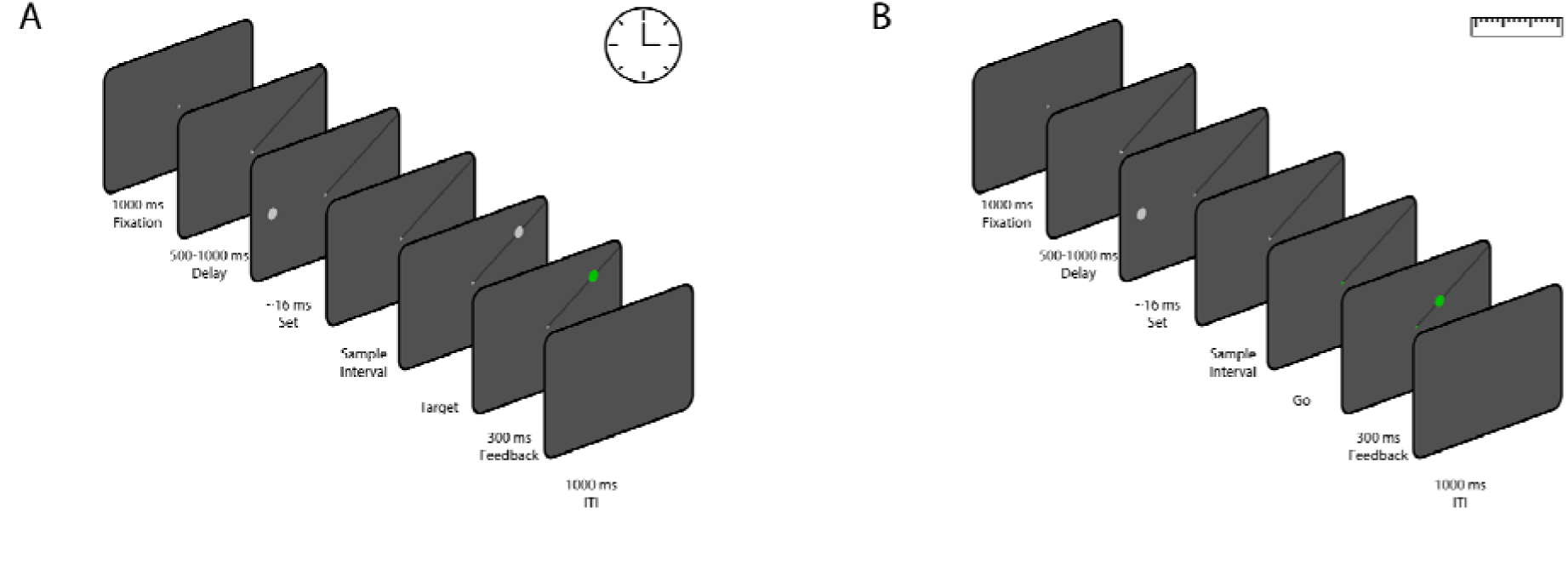
The sequence of trial events for the time reproduction task **(A)** and the distance reproduction task **(B)**. **(A),** Participants were given a time reproduction task in which they had to reproduce sample time intervals. After acquiring fixation and the presentation of a black line on screen, A white circle (‘set’) appeared on the horizontal meridian of the screen, and after a sample interval, another white circle (‘go’) appeared on the black line. Participants had to reproduce the sample interval by making a saccade to the ‘go’ stimulus. Successful saccades turned the ‘go’ stimulus green, but no timing feedback was provided. **(B),** A distance reproduction task was designed similarly to the time reproduction task. It involved the same setup until the appearance of the ‘set’ stimulus. In this case, participants had to reproduce the distance between the ‘set’ stimulus and the fixation cross by making a saccade to a point on the black line at the same eccentricity as the ‘set’ stimulus. Valid saccades were marked with a green circle, and each block consisted of 54 trials with variations in eccentricity and sample intervals.

**Figure 2.**
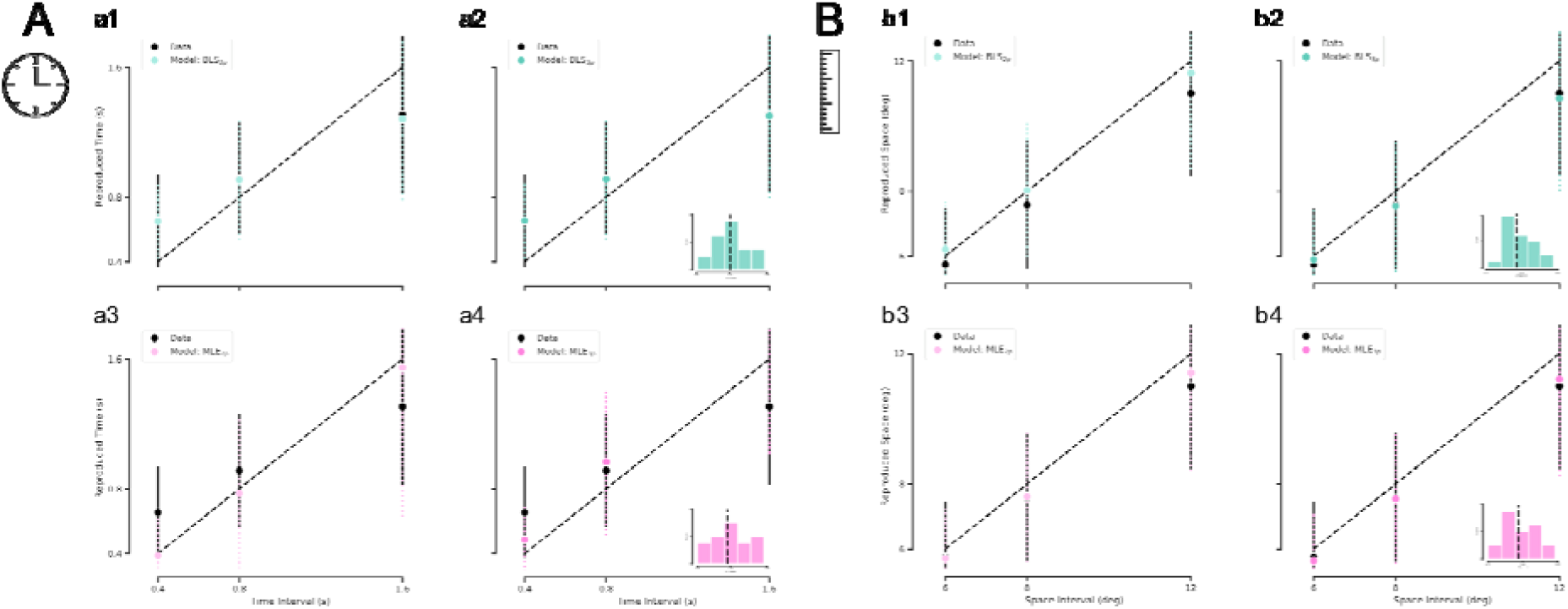
Subjects and observer model behavior in time reproduction **(A)** and distance reproduction tasks **(B)**. **(a1-a4),** Black circles and the error bar show the mean ± SD of subjects’ reproduced times across three sample intervals. The colored circles and the dotted error bar indicate the mean ± SD of the Bayesian Observer model reproduced times computed from simulations of the best-fitted models with 4 estimators (BLS_3p_, BLS_2p_, MLE_3p_, MLE_2p_, respectively). **(b1-b4),** Black circles and the error bar show the mean ± SD of subjects’ reproduced distances across three sample intervals. The colored circles and the dotted error bar indicate the mean ± SD of the Bayesian Observer model reproduced distances computed from simulations of the best-fitted models with 4 estimators, BLS_2p_, BLS_3p_, MLE_2p_, and MLE_3p_, respectively. The inset in **(a2, a4, b2, b4)** shows the distribution of alpha values for all subjects. The dotted line represents the median of alpha distribution.

**Figure 3.**
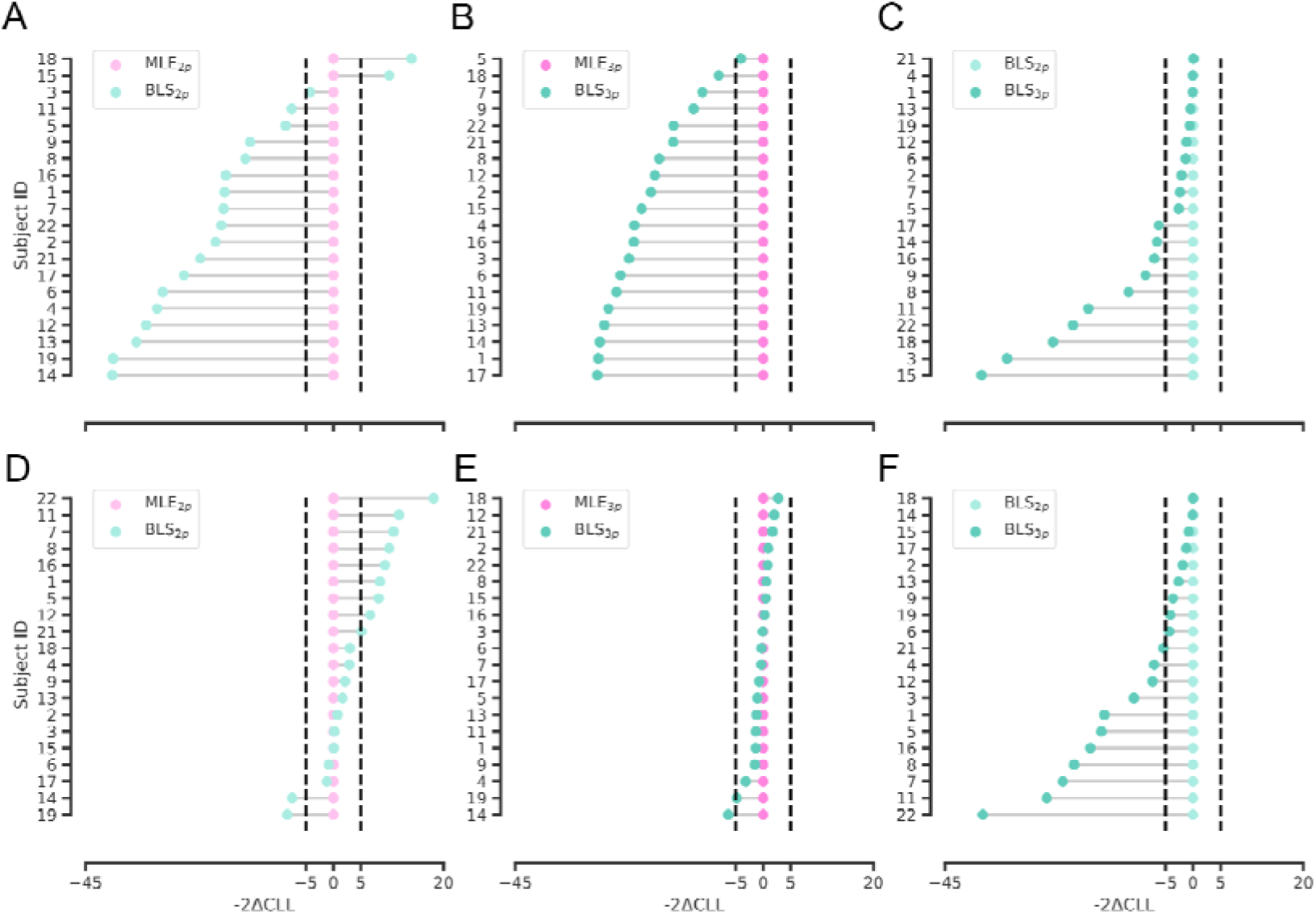
Pair-wise comparisons of observer models with different estimators in time and space. **(A, B)** Cross-validated log-likelihoods (CLL) were multiplied by 2 and the relative differences of BLS_2p_ (light green) and BLS_3p_ (dark green) models to MLE_2p_ (light pink) and MLE_3p_ (dark pink) models in time domain are plotted respectively in **(A)** and **(B)**. **(C)** CLLs were multiplied by 2 and the relative differences of BLS_3p_ (dark green) model to BLS_2p_ (light green) model in time domain is plotted. **(D, E)** CLLs were multiplied by 2 and the relative differences of BLS_2p_ (light green) and BLS_3p_ (dark green) models to MLE_2p_ (light pink) and MLE_3p_ (dark pink) models in space domain are plotted respectively in **(D)** and €. **(F)** CLLs were multiplied by 2 and the relative differences of BLS_3p_ (dark green) model to BLS_2p_ (light green) model in space domain is plotted. Ordinate represents subject’s ID in each subplot.

**Table 1.** Cross-validated log-likelihoods (CLL) computed for models with BLS3p, MLE3p, BLS2p, and MLE2p estimators in *time* domain for all subjects. Bold values represent the highest value among the 4 models.

| Subject ID | BLS<sub>3p</sub> CLL | MLE<sub>3p</sub> CLL | BLS<sub>2p</sub> CLL | MLE<sub>2p</sub> CLL |
| --- | --- | --- | --- | --- |
| 1 | **-3.15** | -18.11 | -3.18 | -13.04 |
| 2 | **-2.25** | -12.43 | -3.29 | -13.98 |
| 3 | **-10.25** | -22.42 | -27.13 | -29.24 |
| 4 | **-12.78** | -24.46 | -12.80 | -28.80 |
| 5 | **-9.62** | -11.66 | -10.92 | -15.25 |
| 6 | **-3.98** | -16.92 | -4.65 | -20.13 |
| 7 | **-9.22** | -14.75 | -10.41 | -20.36 |
| 8 | **-6.99** | -16.43 | -12.85 | -20.84 |
| 9 | **-23.69** | -30.02 | -28.00 | -35.54 |
| 11 | **13.18** | -0.13 | 3.67 | -0.12 |
| 12 | **-21.10** | -30.93 | -21.68 | -38.66 |
| 13 | **-0.74** | -15.14 | -0.99 | -18.87 |
| 14 | **-22.69** | -37.52 | -25.97 | -46.03 |
| 15 | **14.81** | 3.78 | -4.38 | 0.70 |
| 16 | **-11.70** | -23.43 | -15.21 | -24.95 |
| 17 | **-3.50** | -18.55 | -6.61 | -20.16 |
| 18 | **-0.01** | -4.06 | -12.73 | -5.64 |
| 19 | **8.84** | -5.20 | 8.54 | -11.45 |
| 21 | **-16.23** | -24.39 | -16.18 | -28.24 |
| 22 | **-15.24** | -23.37 | -26.14 | -36.31 |

In the distance domain, as it is evident, the 3-paramter models are better fit to the data than the 2-parameter models (Fig. 2B). In 55% of subjects, BLS_3p_ was the best fit among our four models as well (11/20, Table 2). In the pairwise comparisons across models, we observed that in only 2/20 subjects BLS_2p_ was a better fit that MLE_2p_ while in 8/20 subjects, MLE_2p_ was a better fit than BLS_2p_ (Fig. 3D). Comparing BLS_3p_ and MLE_3p_ did not reveal a conclusive picture (Fig. 3E), while BLS_3p_ is a better fit than BLS_2p_ in 11/20 subjects (Fig. 3F). We replicated all of these analyses based on AIC and BIC, and the results were qualitatively similar to CLL data presented here.

**Table 2.**
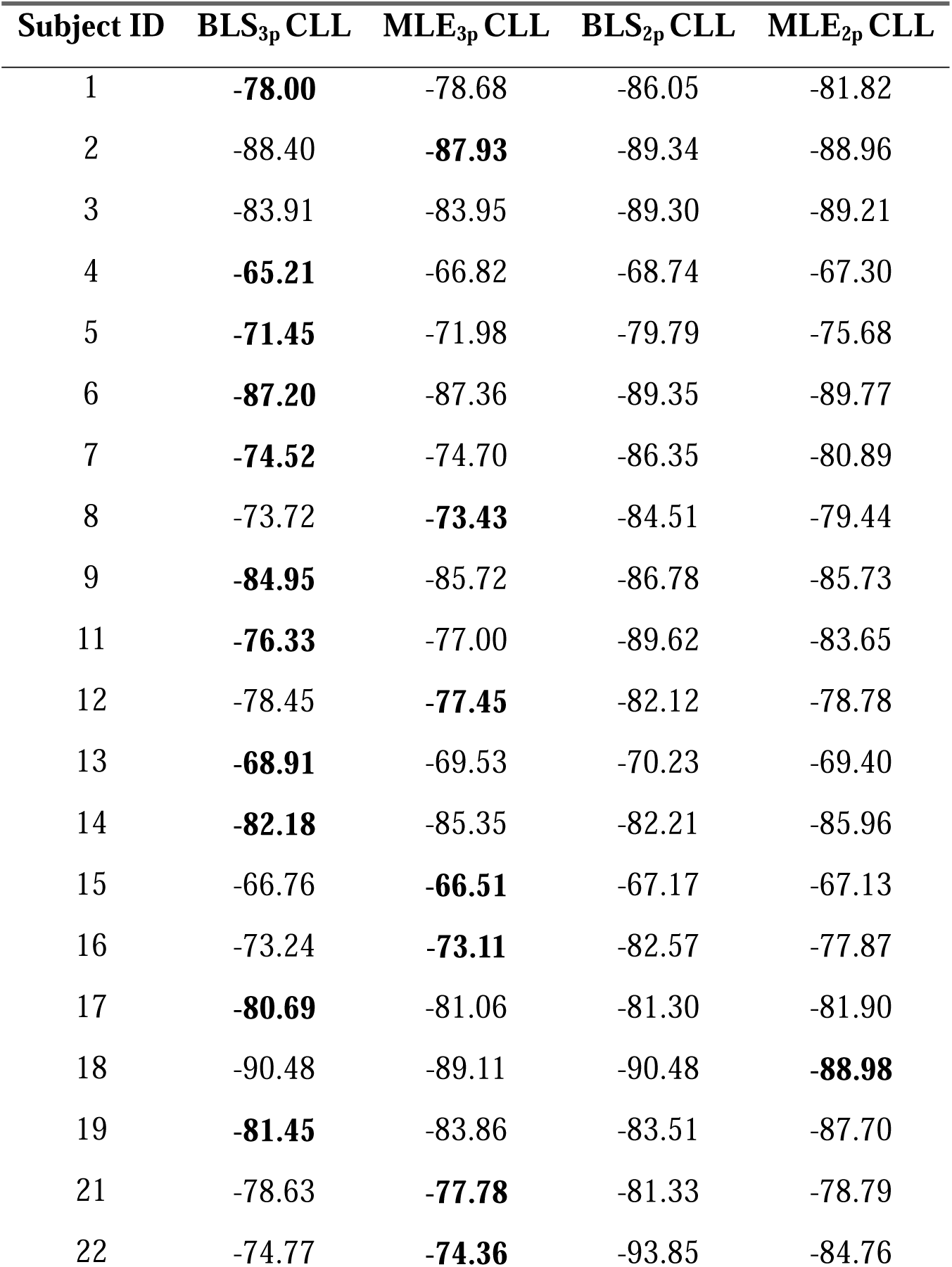
Cross-validated log-likelihoods (CLL) computed for models with BLS3p, MLE3p, BLS2p, and MLE2p estimators in *space* domain for all subjects. Bold values represent the highest value among the 4 models.

### Comparison of model parameters between time and space

Given our results so far, we compared the best fitted free parameters from BLS_3p_ model between time and space. In the case of α we did not find neither correlation (r = -0.11, p = 0.64, Fig. 4A), nor difference between time (mean ± SD: 1.01 ± 0.21) and space (0.93 ± 0.11; W = 70, *p* = 0.2, Wilcoxson Rank Sum). We calculated the measurement (*w_m_*) and the reproduction (*w*_r_) noise parameters of space and time models (Fig. 4B). *w_m_* in space domain (0.05 ± 0.03) was lower than *w_m_* in time domain (0.30 ± 0.07; W = 210, *p* < 0.0001). We also found that *w*_r_ in space domain (0.23 ± 0.02) was smaller than *w*_r_ in time domain (0.29 ± 0.07; W = 185, *p* < 0.001). We found that although there was no correlation between *w_m_* in space and time domain (r = 0.4, *p* = 0.08, Fig. 4C), there was a positive and significant correlation between *w*_r_ in time and space domain (r = 0.45, *p* < 0.05, Fig. 4D).

**Figure 4.**
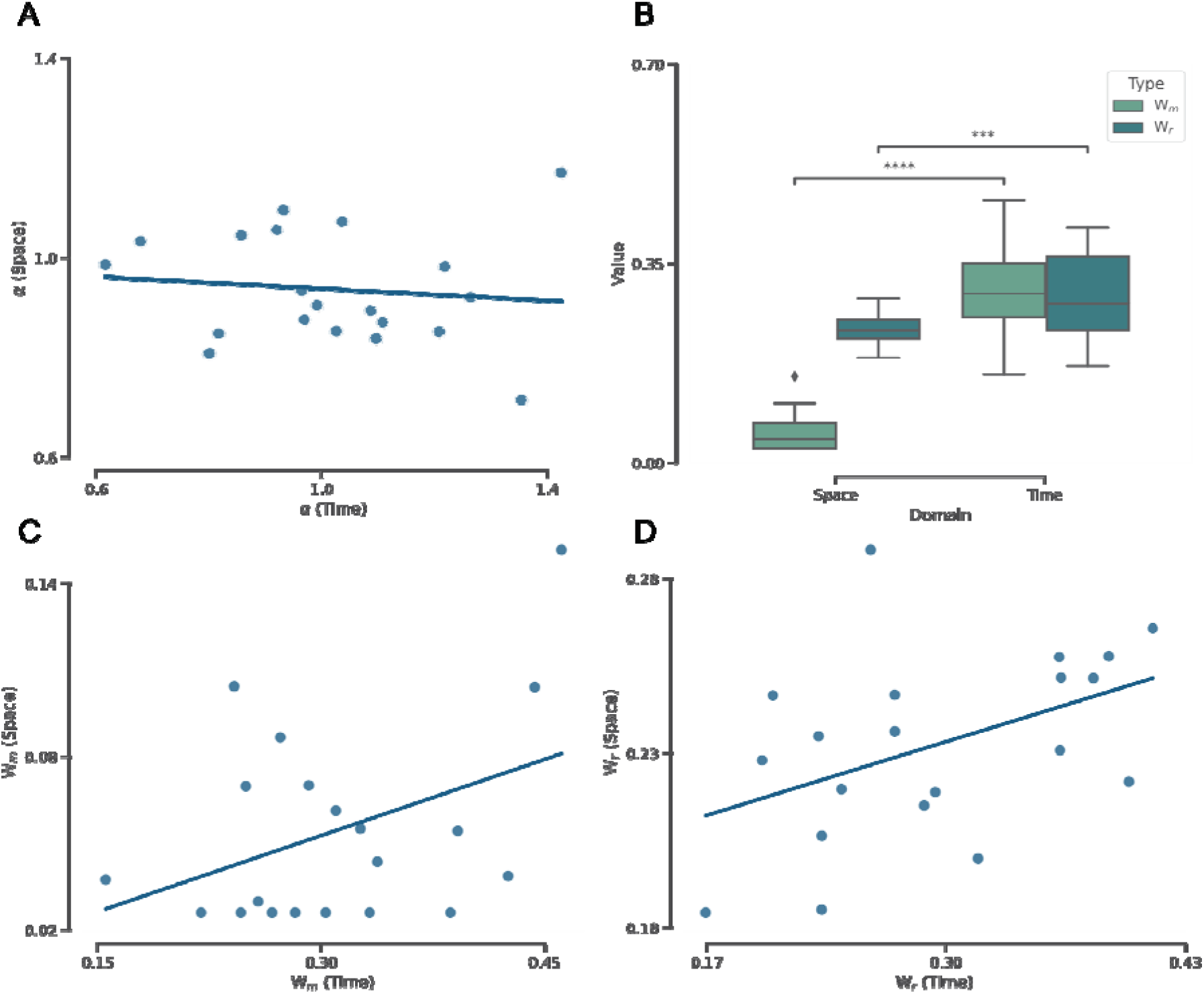
Comparison of and correlation between best fitted model parameters computed in time and space. **(A)** The correlation (r = -0.11, p = 0.64) between alpha computed from the BLS_3p_ model in time and space. **(B)** Box plot comparing the distribution of best-fitted model noise parameters between time and space. The line inside the box represents the median, and the whiskers extend to the most extreme data points within 1.5 times the inter-quartile range (IQR). The Wilcoxon signed rank test was conducted to assess the statistical significance of the differences between best-fitted model noise parameters in time and space. *p* < 0.001***, *p* < 0.0001****. **(C)** The correlation (r = 0.40, p = 0.08) between computed from the BLS_3p_ model in time and space. **(D)** The correlation (r = 0.45, p < 0.05) between computed from the BLS_3p_ model in time and space.

### Prediction of time/distance intervals across different eccentricities/delays

In order to see whether time perception was dependent on distance, we fitted BLS_3p_ on the timing data as a function of stimulus eccentricity and used the fitted parameters for simulation across the three eccentricities (6, 8, and 12°). We computed R^2^ score to assess the goodness-of-fit of a model trained on one eccentricity and tested on other eccentricities. The mean of R^2^ values is shown in Fig. 5A. As can be seen, although each model from any eccentricities can predict data from other eccentricities well (R^2^s > 0.83), data from each eccentricity predicted the same eccentricity better than others (the rightward diagonal in Fig. 5A).

**Figure 5.**
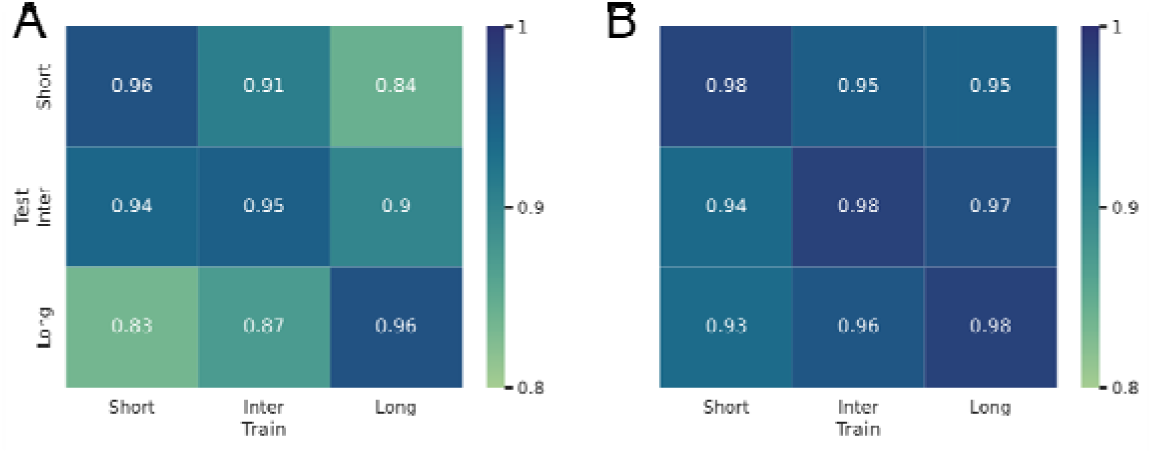
Prediction of a perceptual domain as a function of another one. **(A)** Heatmap shows R^2^ values for the simulated Bayesian Observer model (BLS_3p_) for time domain fitted on trials with short (6°), inter (8°), and long (12°) stimulus eccentricity and tested on trials with these eccentricities. **(B)** Heatmap shows R^2^ values for the simulated Bayesian Observer model (BLS_3p_) for distance fitted on trials with short (0.4 s), inter (0.8 s), and long (1.6 s) fixation to GO delays and tested on trials with these delays.

In order to see whether distance reproduction was dependent on delay interval, we fitted BLS_3p_ on the space data as a function of delay and used the fitted parameters for simulation across the three delays. We computed R^2^ score to assess the goodness-of-fit of a model trained on one delay and tested on other delays (R^2^s > 0.93). The mean of R^2^ values is shown in Fig. 5B. We did not find any systematic difference between models and data as a function of eccentricity.

## Discussion

We investigated the similarities and differences in the perception of time and space, as two components of the ‘A Theory of Magnitude’ (ATOM) proposal ^10^. Our investigation was motivated by the evidence regarding various encoding schemes that are recruited for time and space across different parts of the brain ^19,28,29^. To achieve this goal, we designed two magnitude reproduction tasks and made them as much similar in terms of stimuli and procedure as possible. We used the observer model, which Jazayeri and Shadlen (2010) ^25^ had used, with two different estimators (Bayesian Least Square (BLS) vs. Maximum Likelihood Estimator (MLE)) to our behavioral data. We showed that adding another parameter (α) to the classic Bayesian observer model ^25^ resulted in better fits regardless of the estimator used. We added α to the ideal Observer model in order to capture the heterogeneity of perception of time and space in population ^30^.

We showed that in the time domain, models that take priors into account (i.e., BLS_2p_ and BLS_3p_) outperformed those that do not (i.e., MLE_2p_ and MLE_3p_). In the space domain, it remains inconclusive whether MLE_3p_ or BLS_3p_ better captures our data. Since both of these two models in the space domain fitted data well, we chose BLS_3p_ to compare the time and space with the same model. Given that a probabilistic framework explained both time and space, we believe there is a general computation principle in both domains. Our results are corroborated by studies that have shown an effect of global context in time, but not with those that have shown that in space perception ^22,24,31^. Although time and space perception abide by shared principles, based on the observed differences between the fitted parameters from time and space models, we believe that this probabilistic computation is applied differently in these two domains: time perception is more prior-dependent, while we cannot say the same for space perception.

Our results are supported by biological evidence as well. A recent meta-analysis has shown that although time and space perception share common regions in the brain, their processing might be separated by an anatomical anterior-posterior gradient ^17^. Dissociable neural indices for time and space have also been found in human EEG data ^4^. Neural recordings from prefrontal cortex (PFC) and temporal lobe in epileptic patients found neurons that encode time-only, space-only or both time and space. In non-human primates, researchers found that neurons in the PFC that encode time and space have a small overlap with each other ^29^ and the commonality may be at the level of goal coding. These results can be explained by the recent proposal that the brain uses distinct mechanisms to measure temporal and spatial magnitudes and combines them in a unimodal estimate through another mechanism ^32^.

In the time domain, we observed in subjects with the lowest and the highest α values, α captured the overall underestimation (Fig. S17) and overestimation (Fig. S3) in time reproduction behavior, respectively. We suggest that α represents the speed of an internal clock ^33^. Research has shown that people experience the passage of time in different ways and the speed of the internal clock varies in population ^34,35^. In the space domain, we made the same observation regarding α. However, the majority of subjects had α values under 1 (median of 0.9 for the distribution of α) which translates into an overall underestimation tendency in distance reproduction. This observation is in line with previous report of an average undershoot of about 10% of target eccentricity in peripheral saccades ^36^.

We observed that *w_m_* and *w*_r_, model parameters representing the level of variability, were lower in space than in time models. We also found that *w_m_* was not correlated between time and space explanation for the observed difference in *w_m_* between the two domains could be that there are which hints at the possibility of different measurement mechanisms in the two domains. One various sources of variability for time and space, and the measurement and reproduction of space goes under less noisy stages (or processed further). This difference between time and space can be due to how these dimensions are represented in the brain. We believe that given that brain has many retinotopic maps in which spatial relations such as eccentricity, and distance are coded ^37^, it has the power to reduce noise at the level of measurement. On the other hand, time has very few chronotopic maps ^38^, so the perception of time would be subject to more noise than space. At the motor level, it has long been known that the primate brain has a dedicated system for saccadic eye movement that direct eyes to different locations ^39^. This system is very precise in transforming static visual scenes into spatiotemporal signals for the brain to structure the spatial maps of the environment ^40^. We believe that that is the reason why where to look is less noisy than when to look.

We can compute distance based on information from both egocentric cues (i.e., navigation) and distance from fovea (allocentric, i.e., making a saccade) ^41,42,43^. Distance reproduction tasks so far have mostly focused on egocentric representation as studies have been conducted in virtual reality settings ^44,45^. We used a task design in which the perceived and reproduced distances represent the allocentric mapping of spatial information. In recent years, research has found that there are neurons that encode egocentric spatial representations and also represent allocentric spatial relations in primate hippocampal formation ^46,47^ driven by saccadic eye movements during visual exploration of environment ^48^. Because of this similarity, we believe our results could generalize to tasks that measure the egocentric encoding of distance.

We observed that as the eccentricity of stimulus increased, the subjects overestimated time intervals, as represented in different (not statistically though) α values obtained from models fitted based on eccentricities we had in our task. We also showed that the fitted parameters to timing data in different eccentricities are not generalizable to other eccentricities, as shown in the deterioration of the goodness-of-fit metric. This finding is in line with the previous observations that as the distance between two visual stimuli, with a constant stimulus onset asynchrony, increases, the perceived duration between these two stimuli also increases. This effect is known as Kappa ^8,49^. On the other hand, the reproduced distances as a function of delay were not different. This observation is unexpected given the literature on the relation of working memory and the anti-saccade task which shows the deterioration of task performance as a function of delay interval ^50,51^. We think that the maximum delay duration that we used (1.6 s) is not long enough to manifest the interference effect of time on space.

We do not know that if BLS_3p_ would be a better fit than BLS_2p_ to previously reported data in time domain like the ones from Jazayeri & Shadlen ^25^. But although they trained subjects to have a stable performance before their main task, we did not have that training. They also provided feedback regarding the precision of performance to their subjects on each trial, which we did not have that either. So, although we cannot extend our modeling works to theirs, our model is more powerful in dealing with not giving feedback and also not requiring subjects to be heavily trained. Given that our model can accommodate data collection with minimum training, we believe it can be applied more easily to populations that data collection is an obstacle in them, like children or people with neurological or psychiatric disorders.

## Method

### Apparatus

The experiments were carried out on a computer running Linux operating system, on MATLAB (2016b), with Psychtoolbox 3 extension ^52^. Stimuli were presented on a monitor (17”) placed ∼60 cm from the subject with a 60 Hz refresh rate. The subject sat comfortably on a chair in a dimly lit room to participate in this study, with the head stabilized by a head and chin rest. An EyeLink 1000 infrared eye tracking system (SR Research, Mississauga, Ontario) was used to record eye movements at 1kHz.

### Subjects

We enrolled 22 volunteers (12 female, range: [20, 43], mean ± SD: 26.5 ± 5.5). All were naïve to the purpose of the study except 2 (subjects 1 and 2) who were the authors of this study. We excluded 1 subject because of the troubled eye-tracker calibration caused by her contact lens and 1 subject because of excessively large eye-calibration errors. All subjects had normal or correct-to-normal vision. They had signed the consent form prior to the experiment. The experiment was approved by the ethics committee of the School of Cognitive Sciences (IPM). We counter-balanced all variables and blocks between participants. Half of the participants completed the time reproduction task first. Before starting each experiment, each participant completed a full block of training to familiarize themselves with each task.

### Experiment 1

We designed a time reproduction task in which subjects had to reproduce perceived time intervals. At the beginning of each block, the name of the condition (time) was displayed at the center of the screen. The participant then pressed the space bar to start the block. A white fixation cross with a length of 0.5° was presented at the center of the screen for 1 second. After participants acquired fixation, a black line was then presented for a variable duration of 500-1000ms (uniform distribution). The participants were instructed to keep their gaze on the fixation point (within a 4° × 4° window). The line extended from the fixation cross to one of the four corners of the screen. The location of the line was fixed within each block, but changed between blocks. After that, a white circle (‘set’, diameter of 1.5°) was flashed on the horizontal meridian, contra-lateral to the black line. The eccentricity of the circle was 6, 8, or 12°, randomly chosen on each trial. After a variable sample interval of 0.4, 0.8, or 1.6 seconds, a white circle (‘go’, diameter of 1.5°) was presented on the black line at the same eccentricity as the flashed circle. Participants were then required to reproduce the duration between the onset of the ‘set’ stimulus and the onset of the ‘go’ stimulus by making a saccade to the ‘go’ target to reproduce the sample interval. If the saccade was landed within a 4° × 4° window of the ‘go’ stimulus, the go stimulus would turn to green (Fig. 1A). We did not provide any feedback regarding the accuracy of the timing. Each block consisted of 54 trials (3 eccentricities, 3 sample interval, and 6 repetitions for each condition) and subjects performed 8 blocks.

### Experiment 2

We designed the distance reproduction task as similar to the time reproduction task as possible (Fig. 1B). Each trial is similar to the experiment 1 up to the presentation of the ‘set’ stimulus. Here, after passing a variable duration (0.4, 0.8, or 1.6 seconds) from ‘set’ stimulus onset, the fixation cross turned to green (go signal). This indicated to the participants that they should reproduce the distance between the ‘set’ stimulus and the fixation cross by making a saccade to a point on the black line that had the same eccentricity as the ‘set’ stimulus. Gaze locations which landed within 2° of the black line were considered valid. A green circle (diameter of 1.5°) was presented at the location of the saccade. Each block consisted of 54 trials (3 eccentricities, 3 sample interval, and 6 repetitions for each condition) and subjects performed 8 blocks.

### Data Analysis

We employed the Interquartile Range (IQR) method as a robust statistical technique to identify and eliminate outlier data points ^53^. This method excludes the data that lie outside of 1.5 IQR below the first quantile or 1.5 IQR above the third quantile. We applied the IQR method for each subject for each experiment. We also excluded trials with a reaction time of less than 200 ms The number of excluded trials per subject per experiment was below 1%.

### Ideal Observer model

In our data, we had pairs of sample time/distance intervals (*t_s_, t_s_*) and corresponding reproduced times/distances (*t_r_, t_r_*) for each trial. We used an ideal observer model to relate sample times/distances to the reproduced ones. To model these relationships, we used two hidden variables, each of which refers to one of the noisy stages of the process of reproducing time/distance intervals. The model has three stages as:

Measurement stage

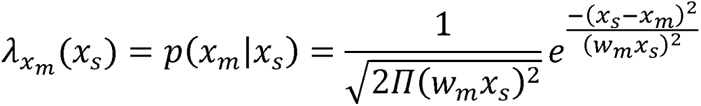

Estimation stage

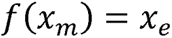

Reproduction stage

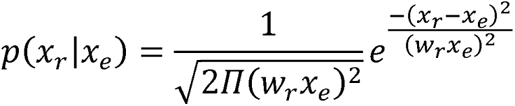

and then

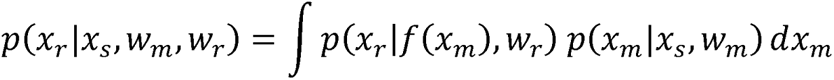

x stands for t (parameters from the time reproduction task) or d (parameters from the space reproduction task)

For the time reproduction task, we have *t_s_*, sample time interval; *t_m_*, measured time interval; *t_r_*, reproduced time interval; *t_e_*, estimated time interval; *w_m_*, measurement Weber fraction; *w*_r_, reproduction weber fraction. For the distance reproduction task, we have *d_s_* sample distance interval, *d_m_*, measured distance interval, *d_r_*, reproduced distance interval, and, *d_e_*, estimated distance interval.

In these observer models, *p*(*t_m_*|*t_s_*) and *p*(*d_m_*|*d_s_*) are modeled as Gaussian distributions centered at *t_s_* and *d*_s_, and we assume that their standard deviations (SD) grow linearly with their means. This assumption is motivated by the scalar variability of timing and distance ^25,27^. The of *p*(*t_m_*|*t_s_*) and *p*(*d_m_*|*d_s_*), which we will refer to as the Weber fraction associated with the distribution of measurement noise is thus fully characterized by the ratio of the SD to the mean measurement, *w_m_*. With the same arguments in mind, we assume that the distributions of *t_r_* and *d_r_* conditioned on *t_e_* and *d_e_*, *p*(*t_r_*|*t_e_*) and *p*(*d_r_*|*d_e_*), are also Gaussian, centered at *t_e_* and *d_e_*, and associated with a constant Weber fraction, *w_r_*. We used a Maximum Likelihood Estimation (MLE) function, which does not fuse prior information with the likelihood function, and a Bayesian Least Squares (BLS) function in the estimation stage of both tasks. For the Bayesian models, we used a uniform distribution over the range of experimental 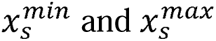. The BLS and MLE functions were defined as:

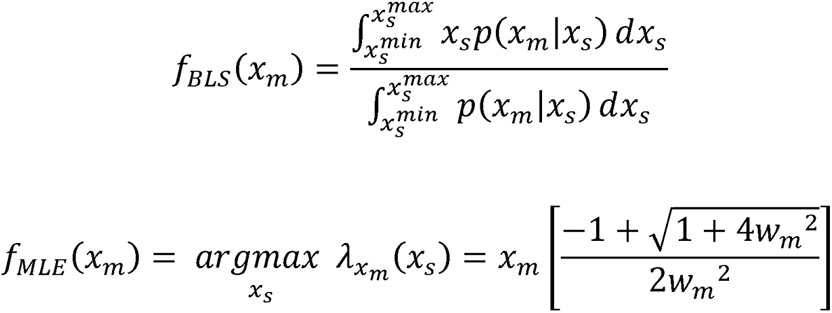

In the pilot data, we observed individual-specific shifts in the range effect as represented in an overall tendency to over/under-estimate across the whole range of intervals. These patterns were not explained by the common observer model so we used a modified version of these models. We introduced another free parameter (α) to the estimation stage of the model as a multiplication factor. So, the estimation stage for these models would be:

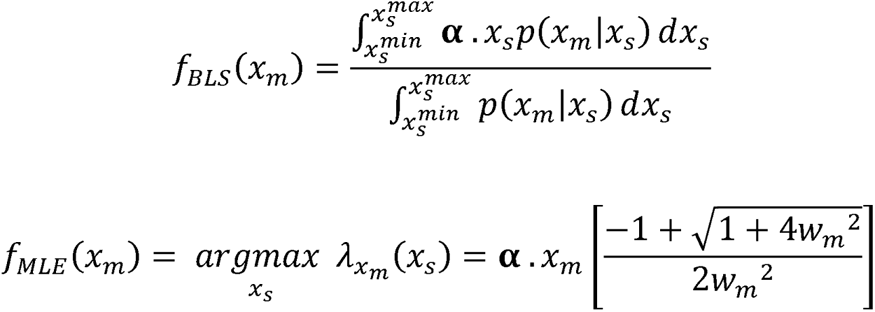

In a separate analysis, we added α to the estimation stage as an additive parameter. The result from multiplication and addition did not differ from each other qualitatively, so we only showed the multiplicative modulation. We preferred the multiplication result as it gives us a unitless α which is comparable between our tasks.

### Model fitting

We maximized the likelihood of model parameters w*_m_*, w*_r_*, and α (when applicable) across all *x_s_* and *x_r_* values. Maximum likelihood estimation was performed with the minimize function in SciPy library, using the Nelder-Mead downhill simplex optimization method. We evaluated the success of the fitting procedure by repeating the search with several different initial values.

### Model comparison

In order to compare modified models (with α) with the previous models (without α), we used Akaike Information Criterion (AIC), Bayesian Information Criterion (BIC), and Cross-validated Log-Likelihoods (CLL) as our quantitative criteria. We also plotted the averaged result of 50 simulations of tasks with the best fitted models over the data. We compared plots to make sure there are visible differences between the models. We also wanted to check that the best model actually captures the pattern of the data well. We considered differences bigger than 5 in each criterion (i.e., AIC, BIC, and CLL) between different models, as an indicator of better model performance ^54^. We also used the highest CLL values among the models as an absolute measure of goodness-of-fit. We performed comparisons between the modified and the classic models, separately for f_BLS_ and f_MLE_ estimators, to choose the best model for time and space reproduction, again separately.

### Difference between model parameters of time and space

We employed Wilcoxon signed-rank test to detect possible differences between model parameters (α, *w_m_*, and *w_r_*) obtained from the best fitted BLS_3p_ model between time and space. We considered p-values of less than 0.05 as an indication of statistical significance.

### Correlation between time and space

We calculated Pearson correlation between the best fitted model parameters for the time and distance reproduction tasks to measure the degree of potential overlap between time and distance perception.

### Prediction of a perceptual domain as a function of another one

We fitted BLS_3p_ on the time and distance data as a function stimulus eccentricity and fixation to GO delay, respectively. We computed R^2^ score to assess the goodness-of-fit of a model trained on one eccentricity/delay and tested on other eccentricities/delays. Since we wanted to see how the overall performance of our model in capturing the mean and SD of data across different eccentricities/delays changes, we used the mean predicted values and the true values across time and distance intervals to calculate R^2^ for each subject. With this approach we had the problem of small number of sample points (3 in each domain) which resulted in negative R^2^ values in some of the train-test combinations in time domain for 5 subjects. We excluded these 5 subjects from this analysis in time domain.

### Data Availability

Data and code will be published upon acceptance of the paper.

## Acknowledgments

We are grateful to Abdol-hossein Vahabie for giving feedback on our data analysis approach, and Mohammad Rabie for comments on a previous version of our manuscript.

---

**Figure S1-S20.** Subject and observer model behavior in time reproduction (Up) and in distance reproduction (Down).

