## Supplementary material for "Same principle, but different computations in representing time and space": Supplemntal figures

### 1 Supplementary Figures and Tables

#### 1.1 Supplementary Figures

**Figure S1.** Subject and observer model behavior in time reproduction (Up) and in distance reproduction (Down).
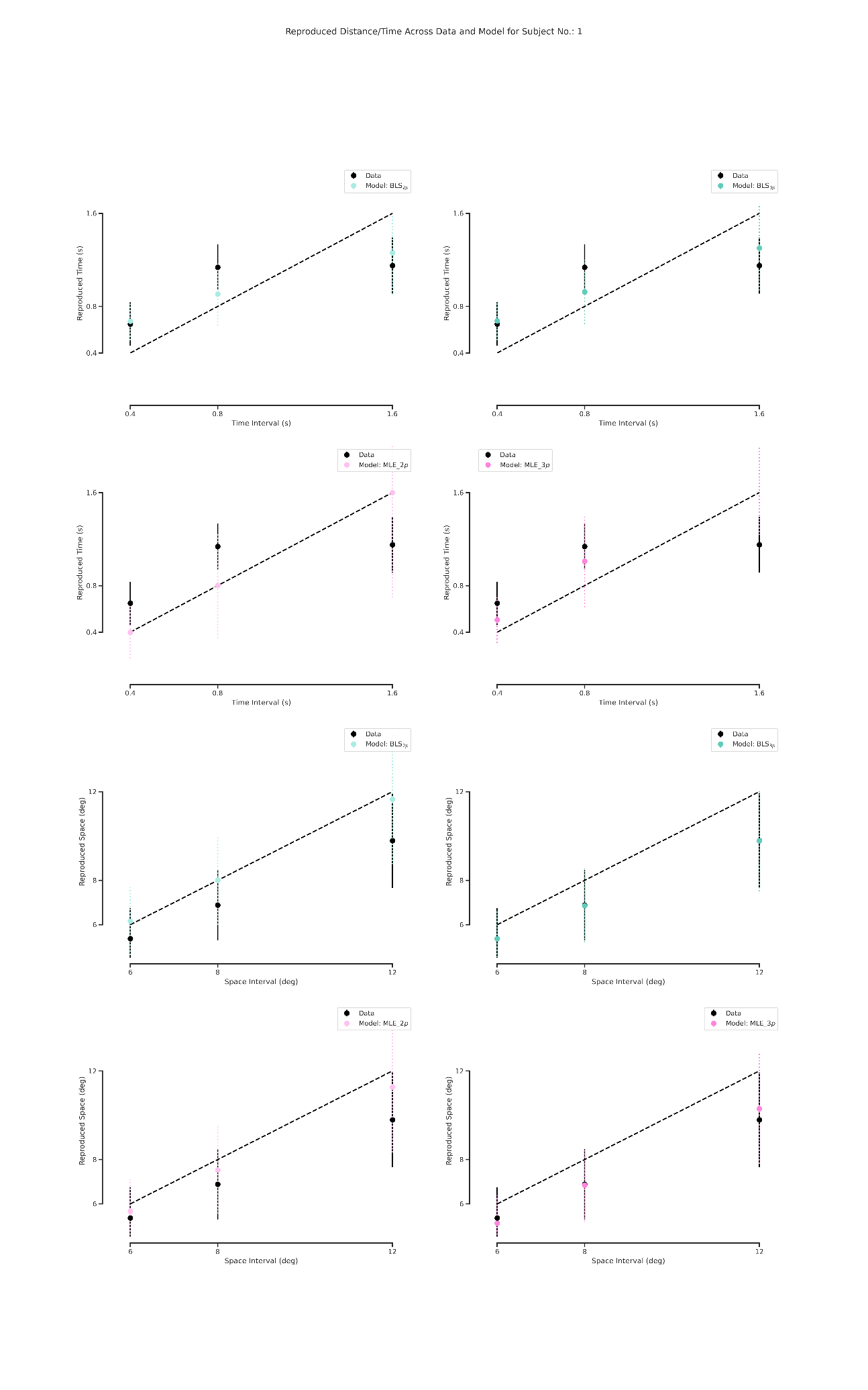

**Figure S2.** Subject and observer model behavior in time reproduction (Up) and in distance reproduction (Down).
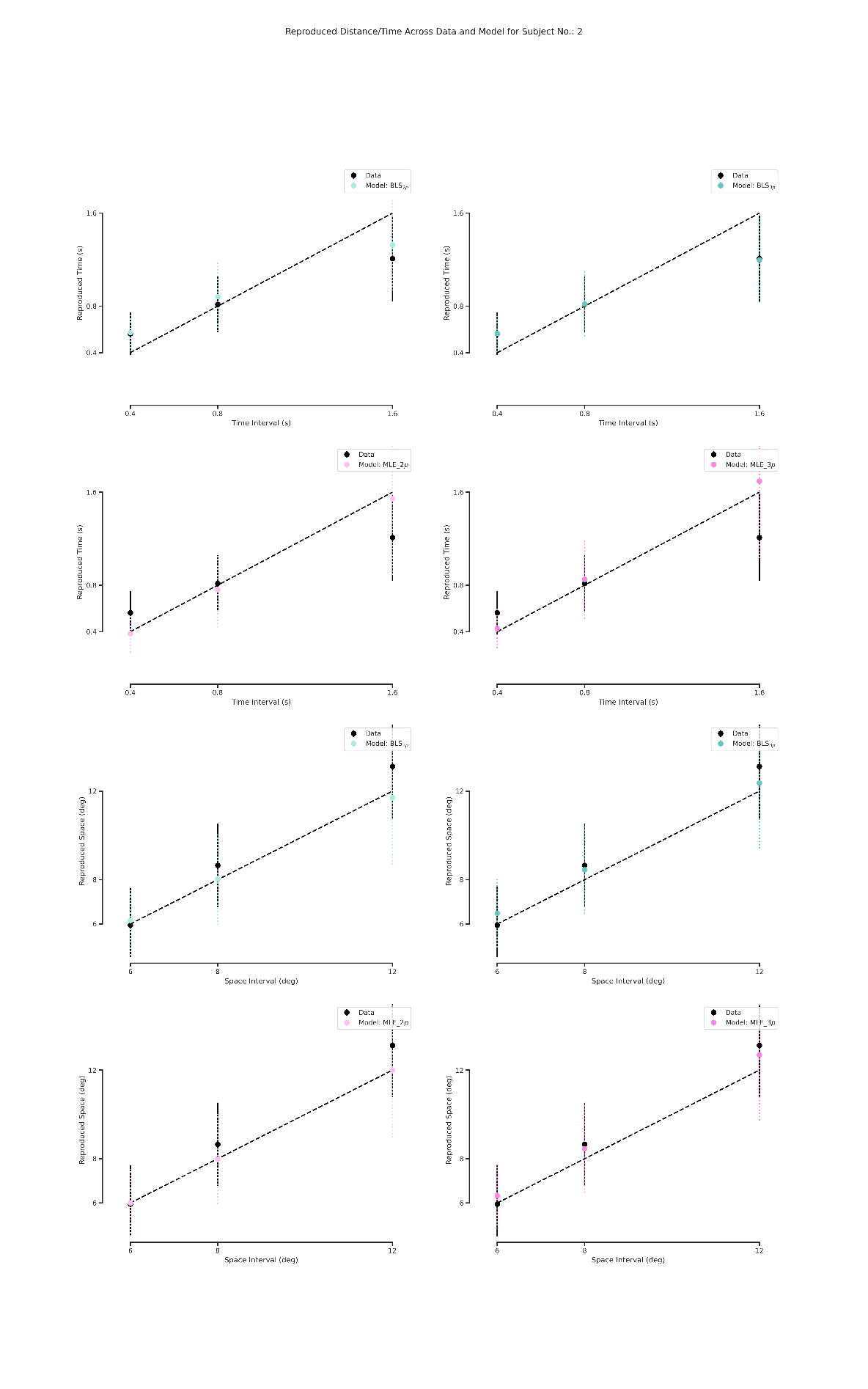

**Figure S3.** Subject and observer model behavior in time reproduction (Up) and in distance reproduction (Down).
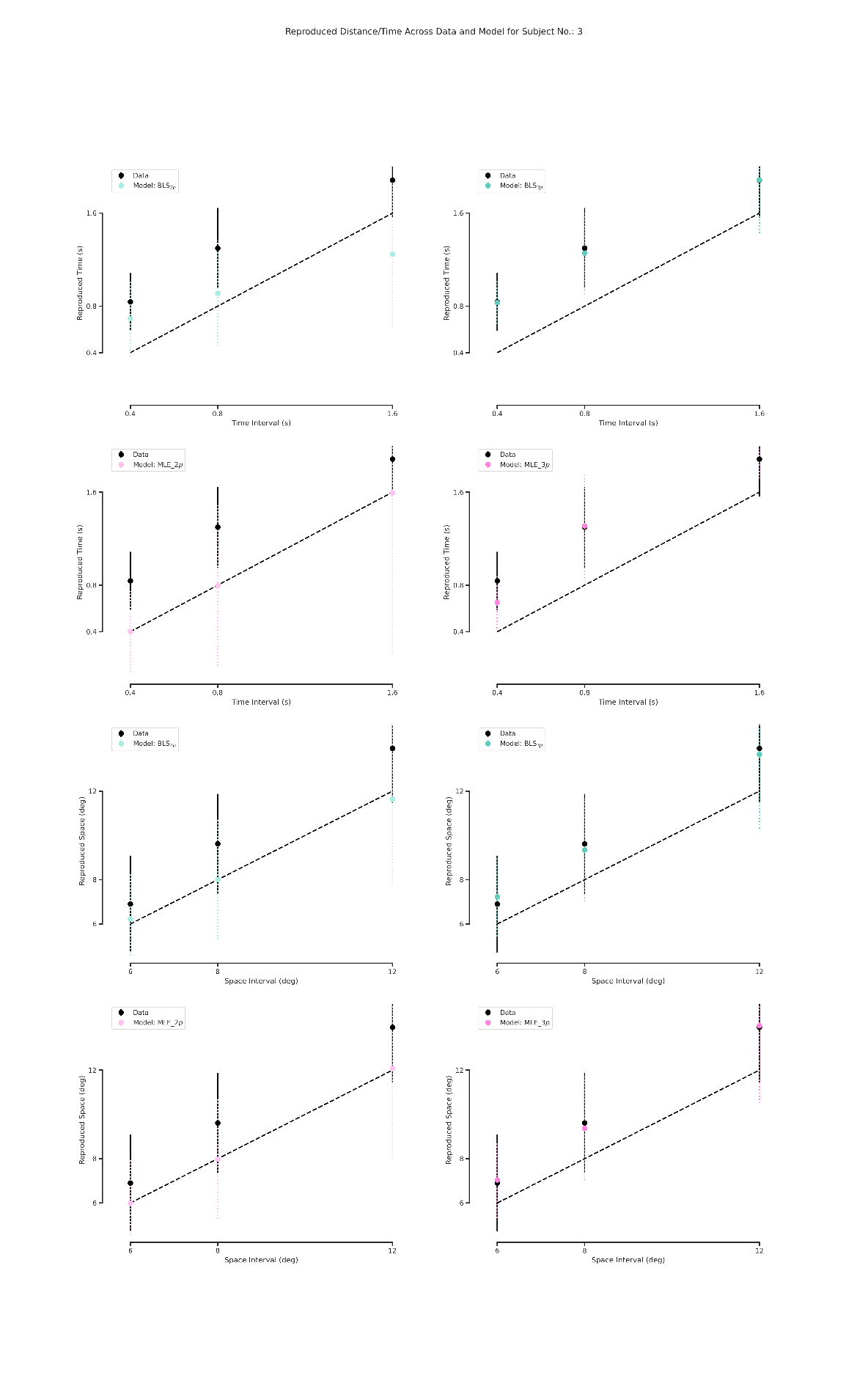

**Figure S4.** Subject and observer model behavior in time reproduction (Up) and in distance reproduction (Down).
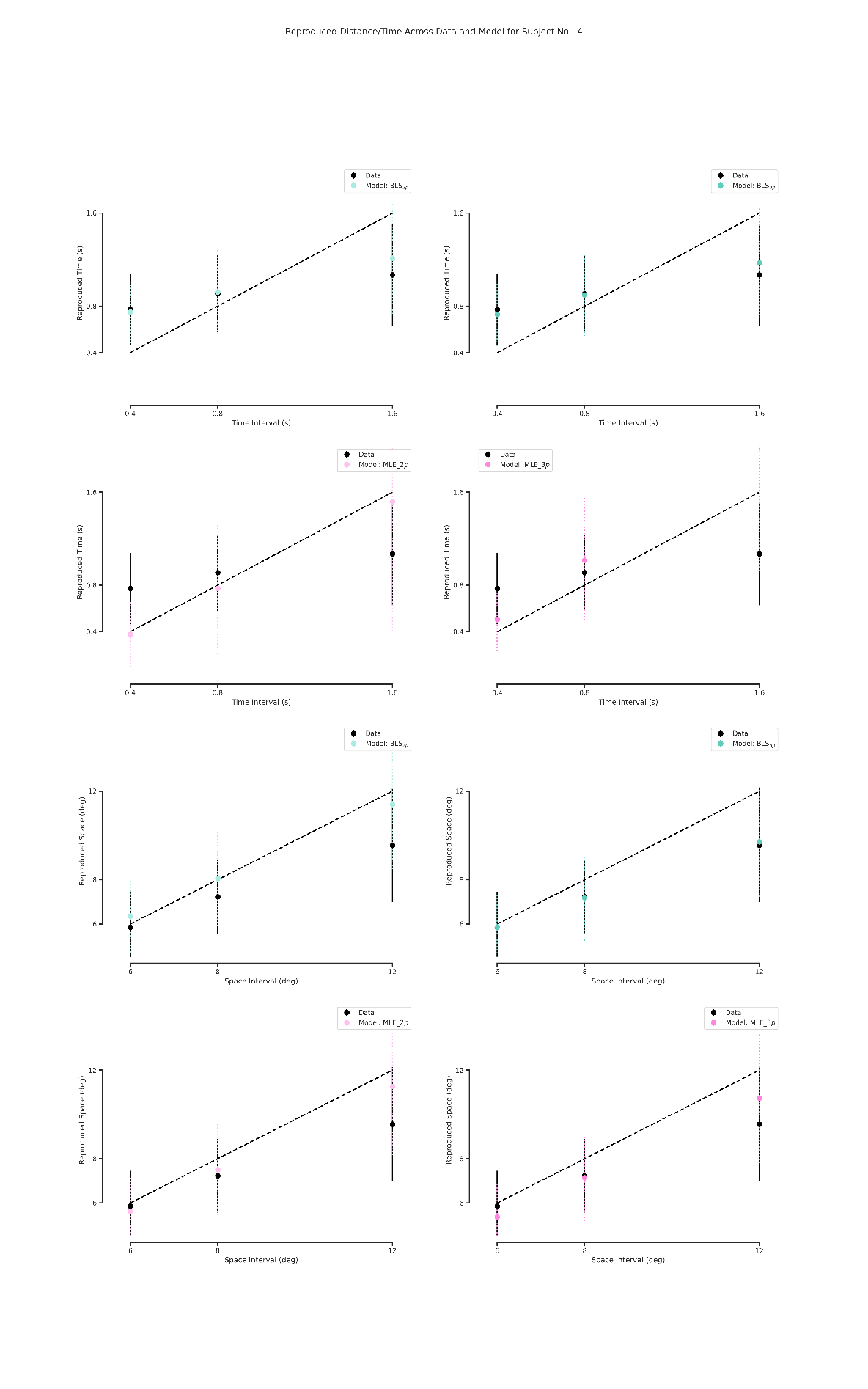

**Figure S5.** Subject and observer model behavior in time reproduction (Up) and in distance reproduction (Down).

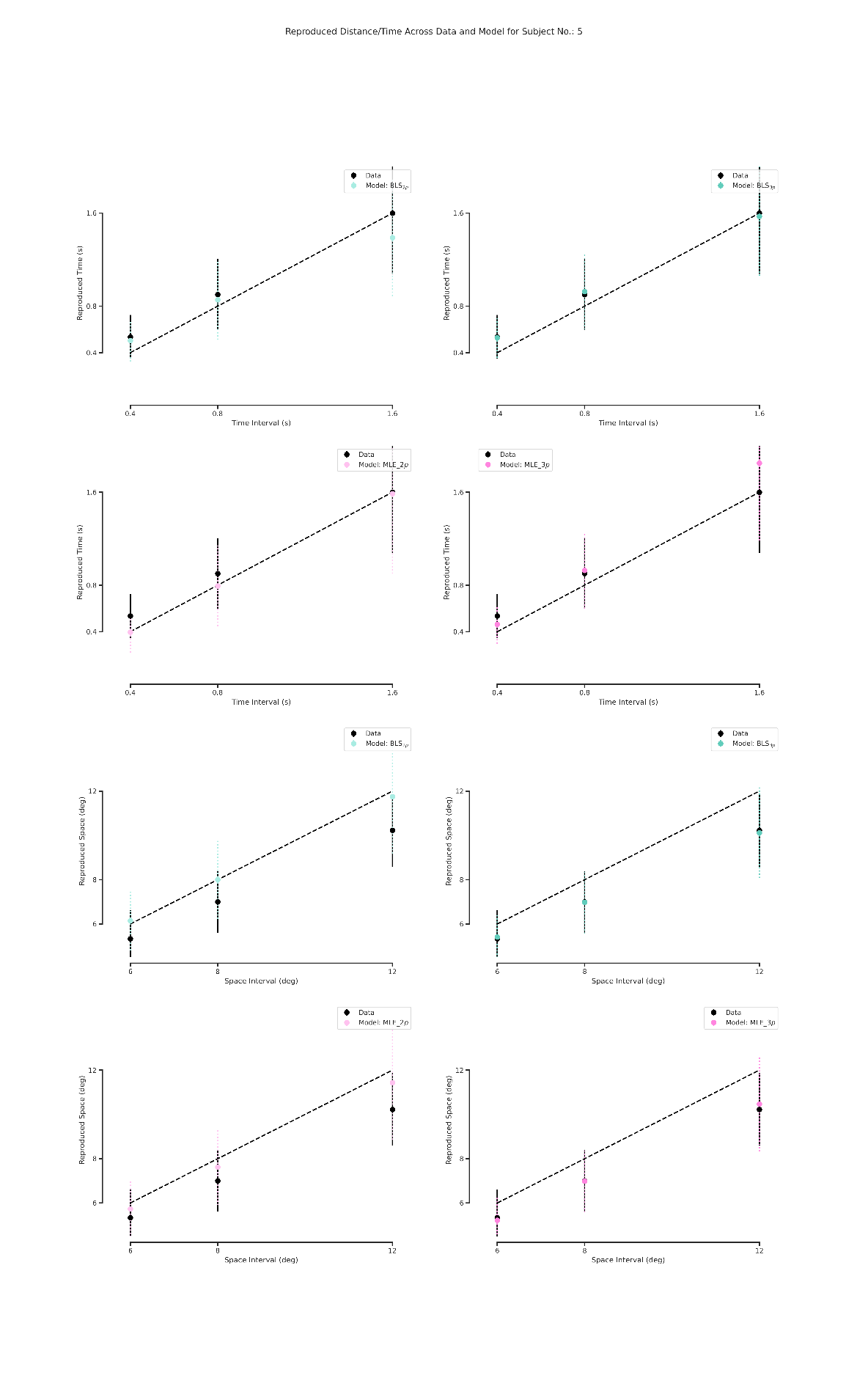
**Figure S6.** Subject and observer model behavior in time reproduction (Up) and in distance reproduction (Down).
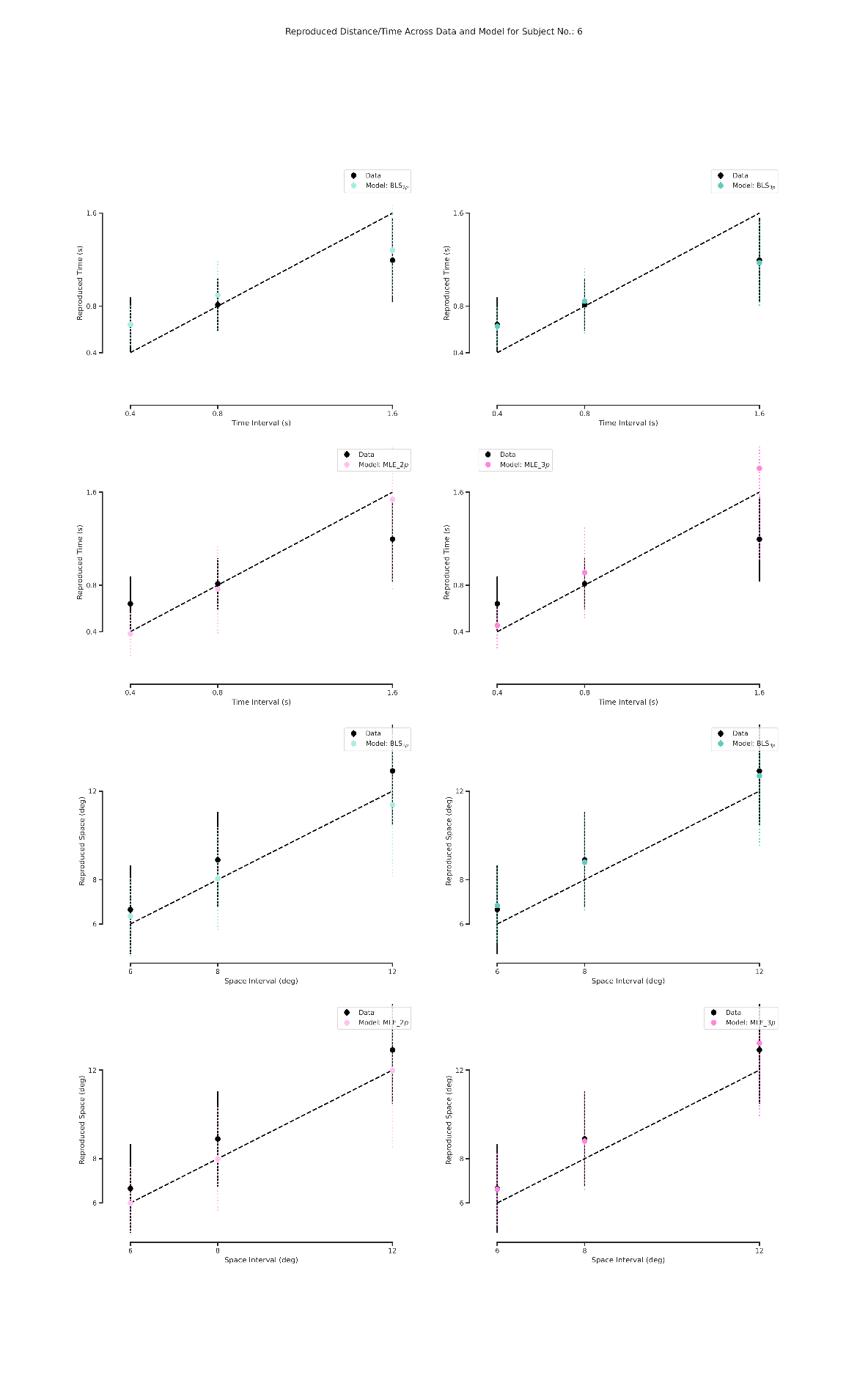

**Figure S7.** Subject and observer model behavior in time reproduction (Up) and in distance reproduction (Down).
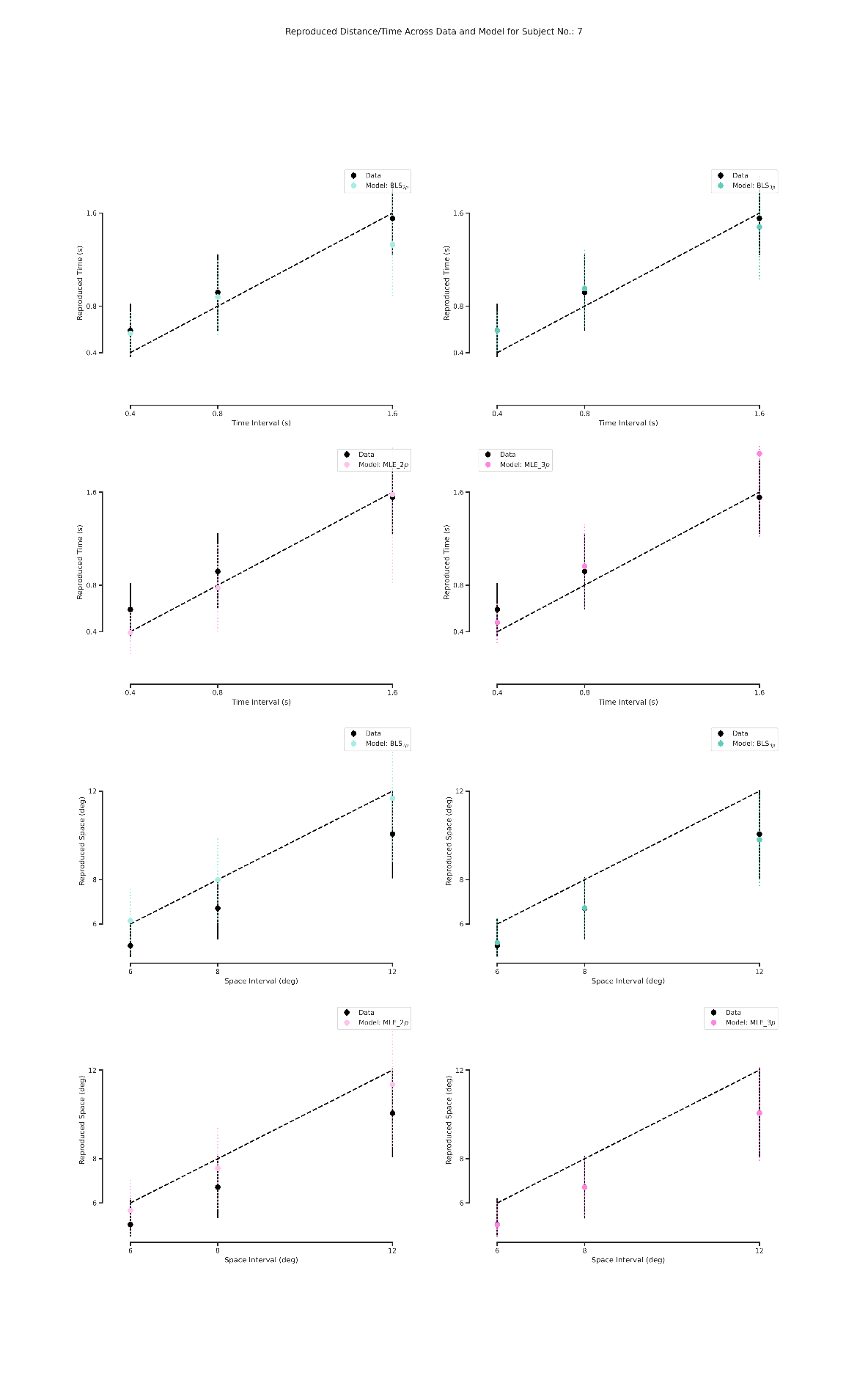

**Figure S8.** Subject and observer model behavior in time reproduction (Up) and in distance reproduction (Down).
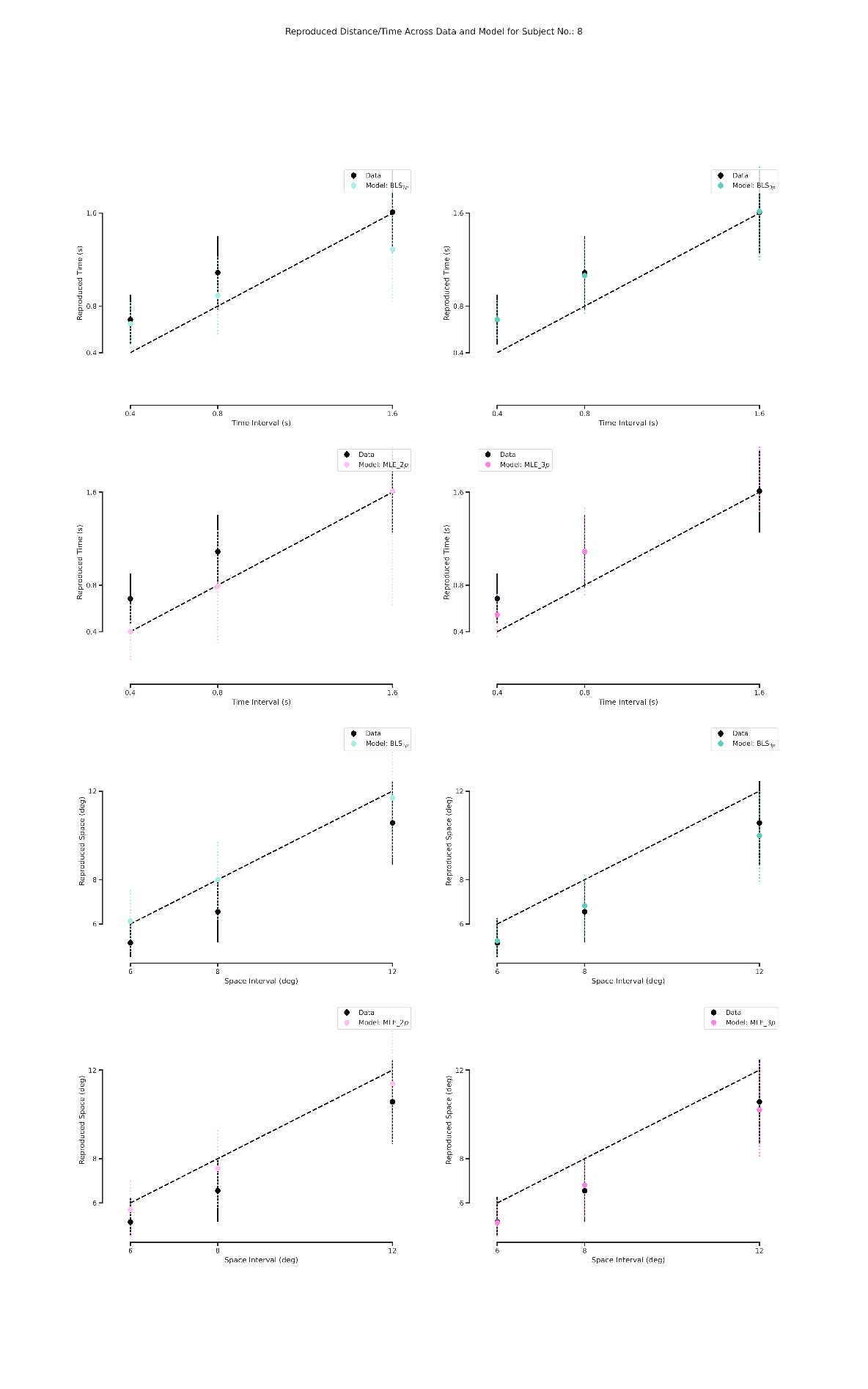

**Figure S9.** Subject and observer model behavior in time reproduction (Up) and in distance reproduction (Down).
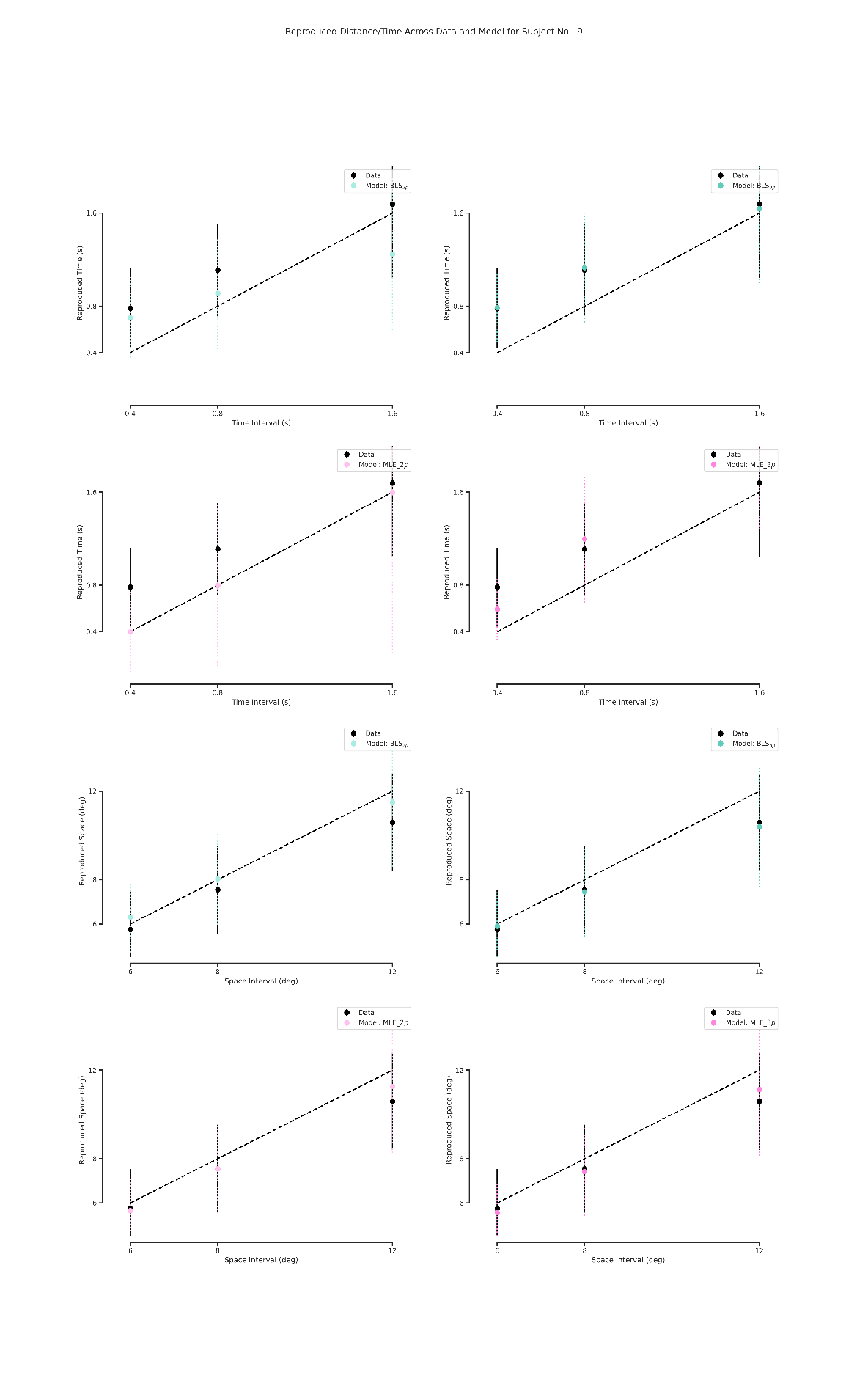

**Figure S10.** Subject and observer model behavior in time reproduction (Up) and in distance reproduction (Down).
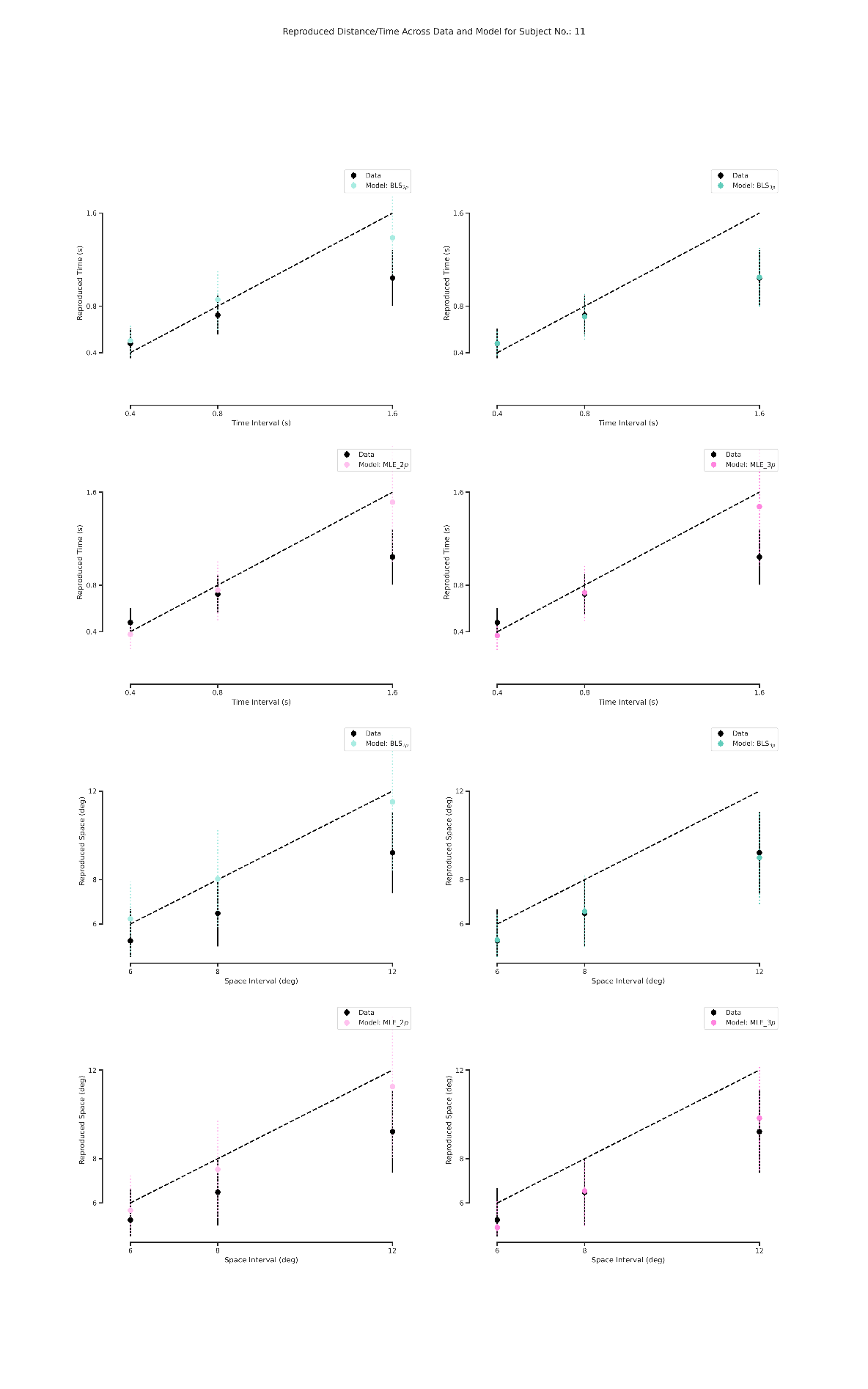

**Figure S11.** Subject and observer model behavior in time reproduction (Up) and in distance reproduction (Down).

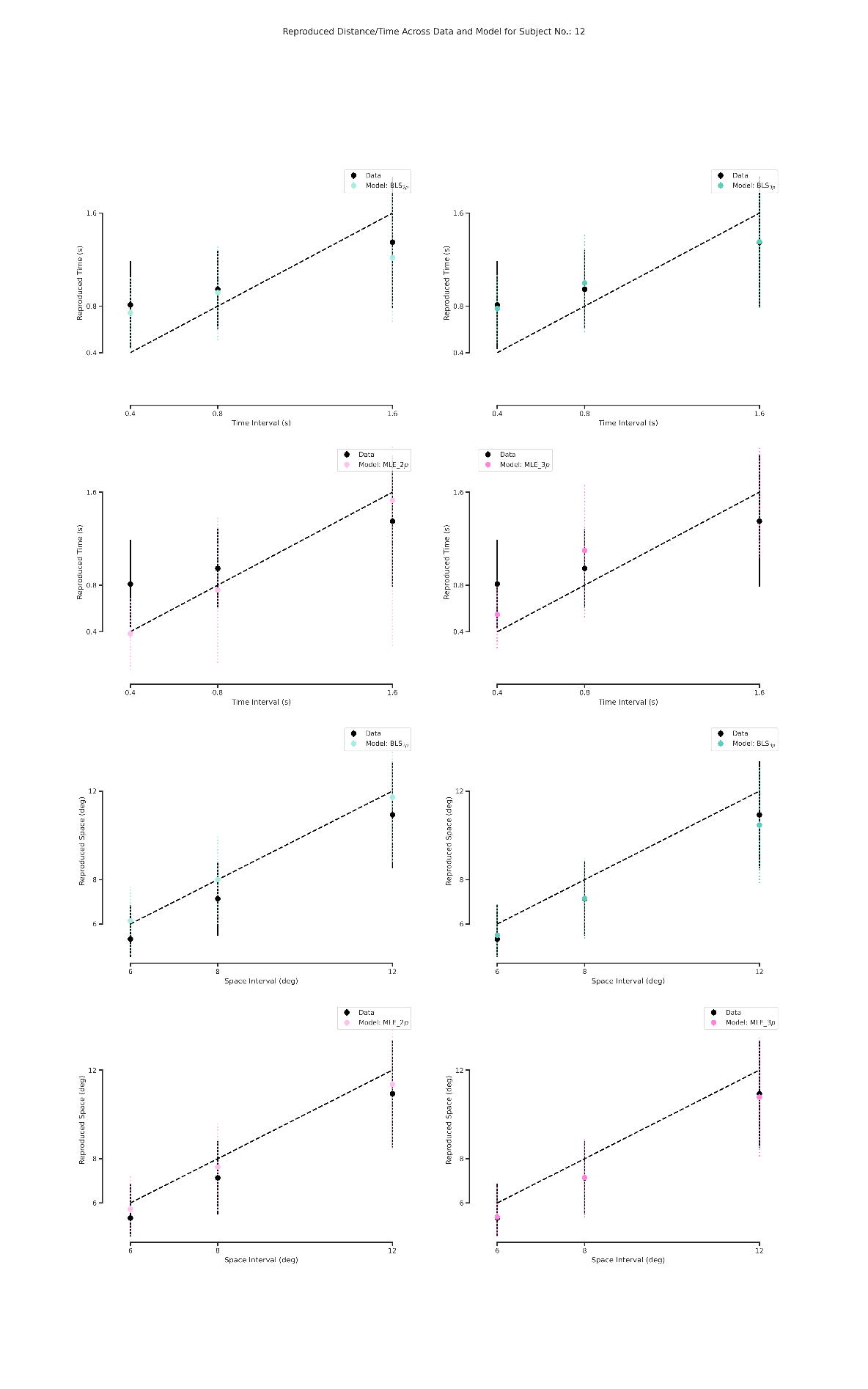

**Figure S12.** Subject and observer model behavior in time reproduction (Up) and in distance reproduction (Down).
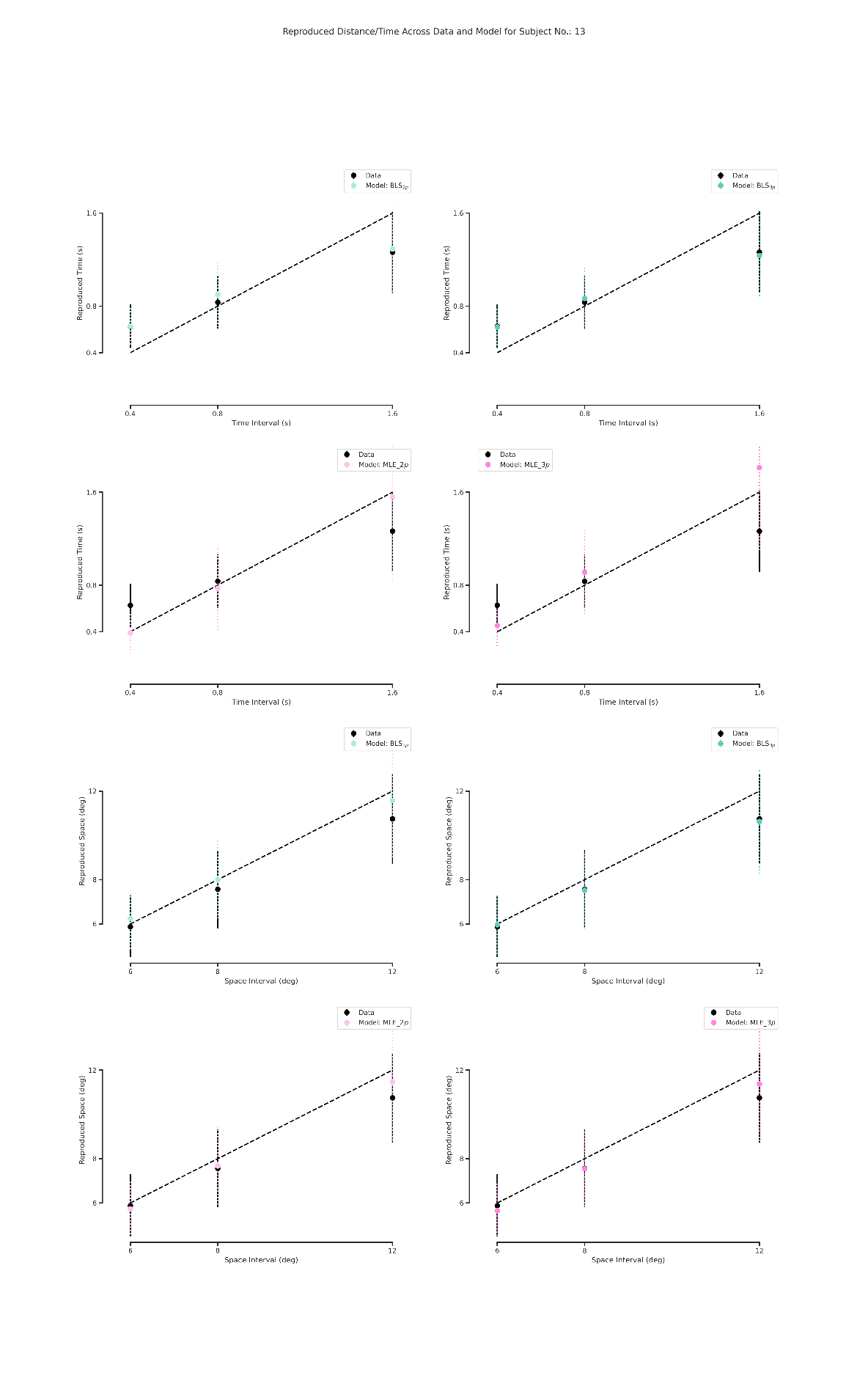

**Figure S13.** Subject and observer model behavior in time reproduction (Up) and in distance reproduction (Down).

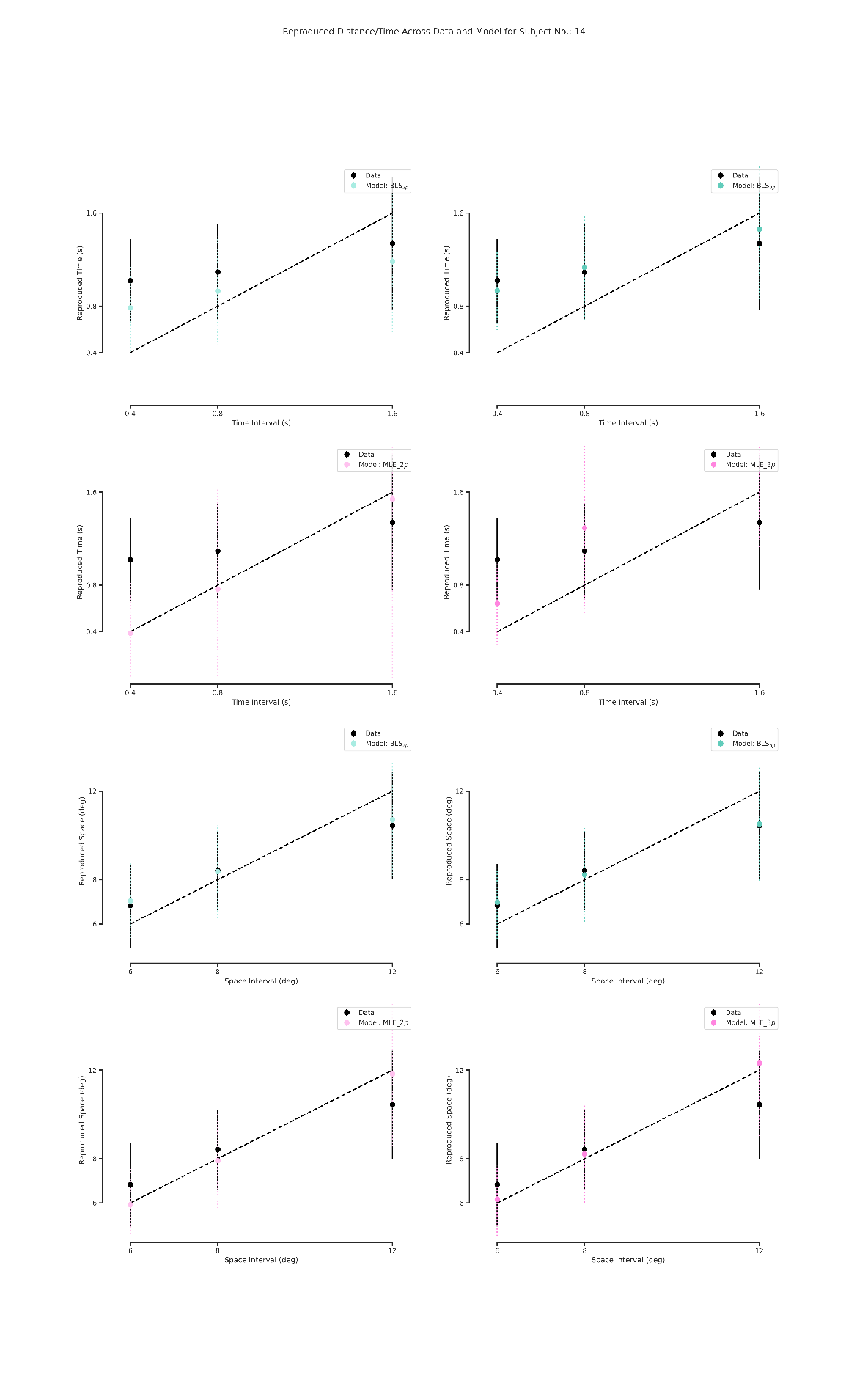

**Figure S14.** Subject and observer model behavior in time reproduction (Up) and in distance reproduction (Down).
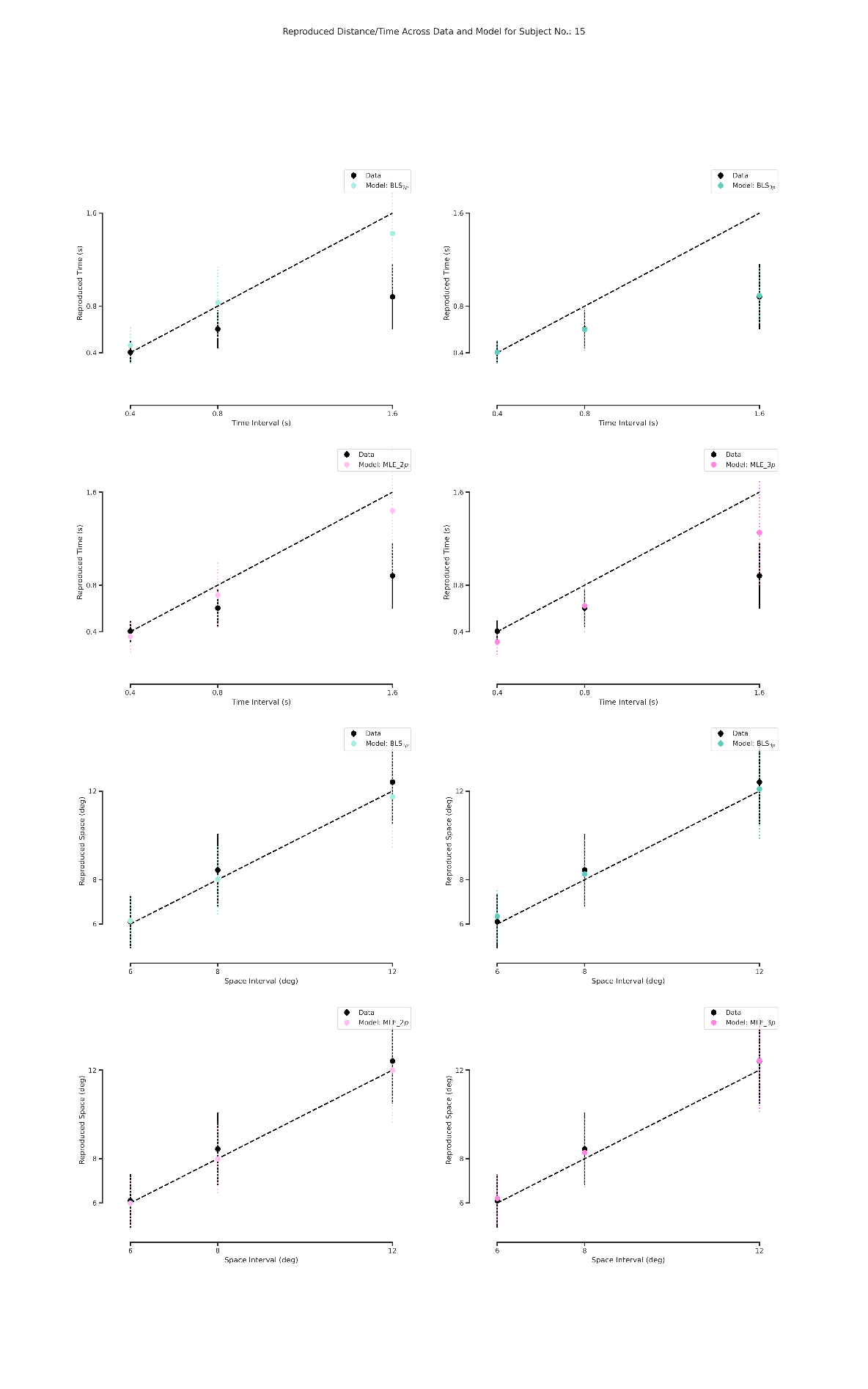

**Figure S15.** Subject and observer model behavior in time reproduction (Up) and in distance reproduction (Down).

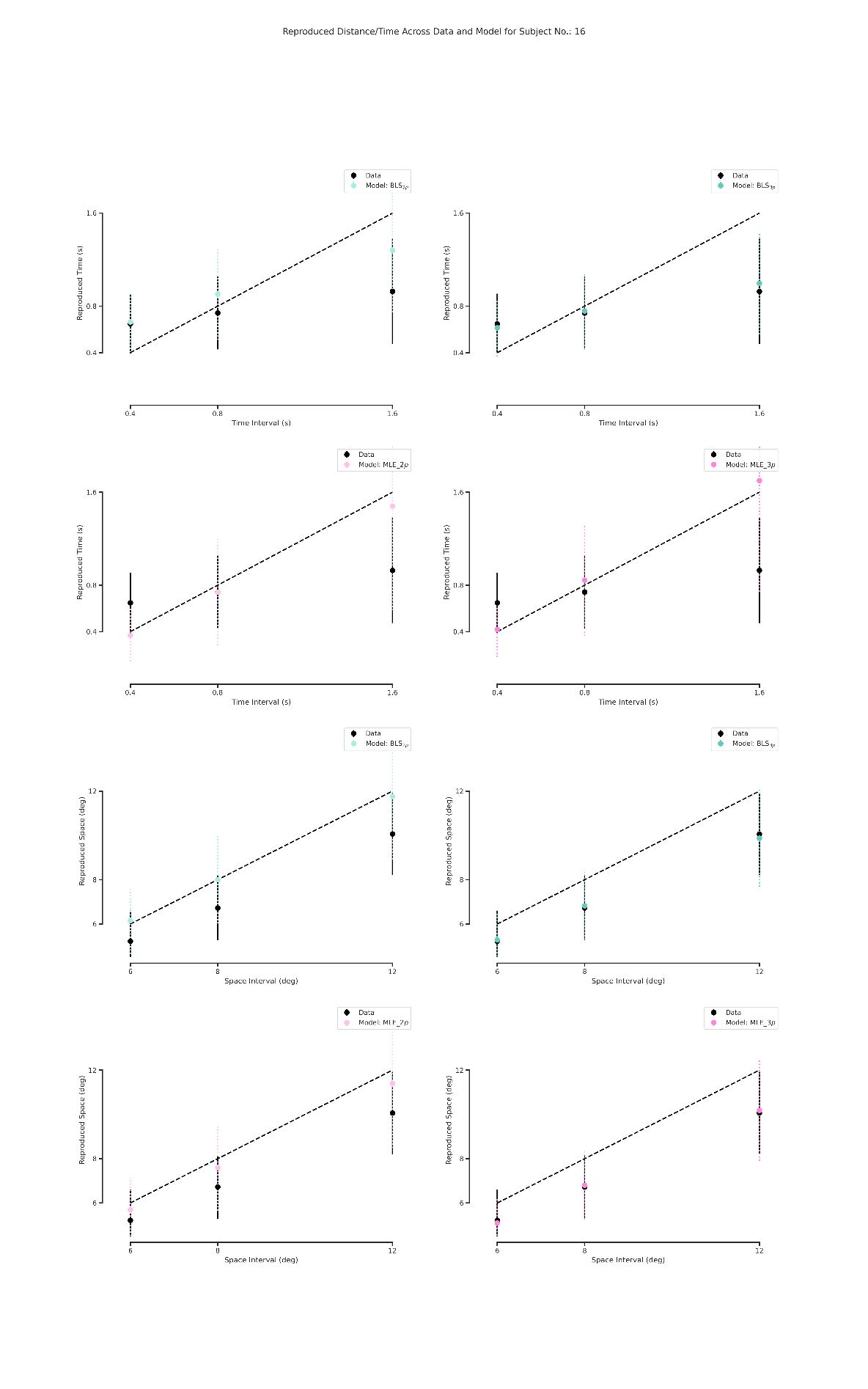

**Figure S16.** Subject and observer model behavior in time reproduction (Up) and in distance reproduction (Down).
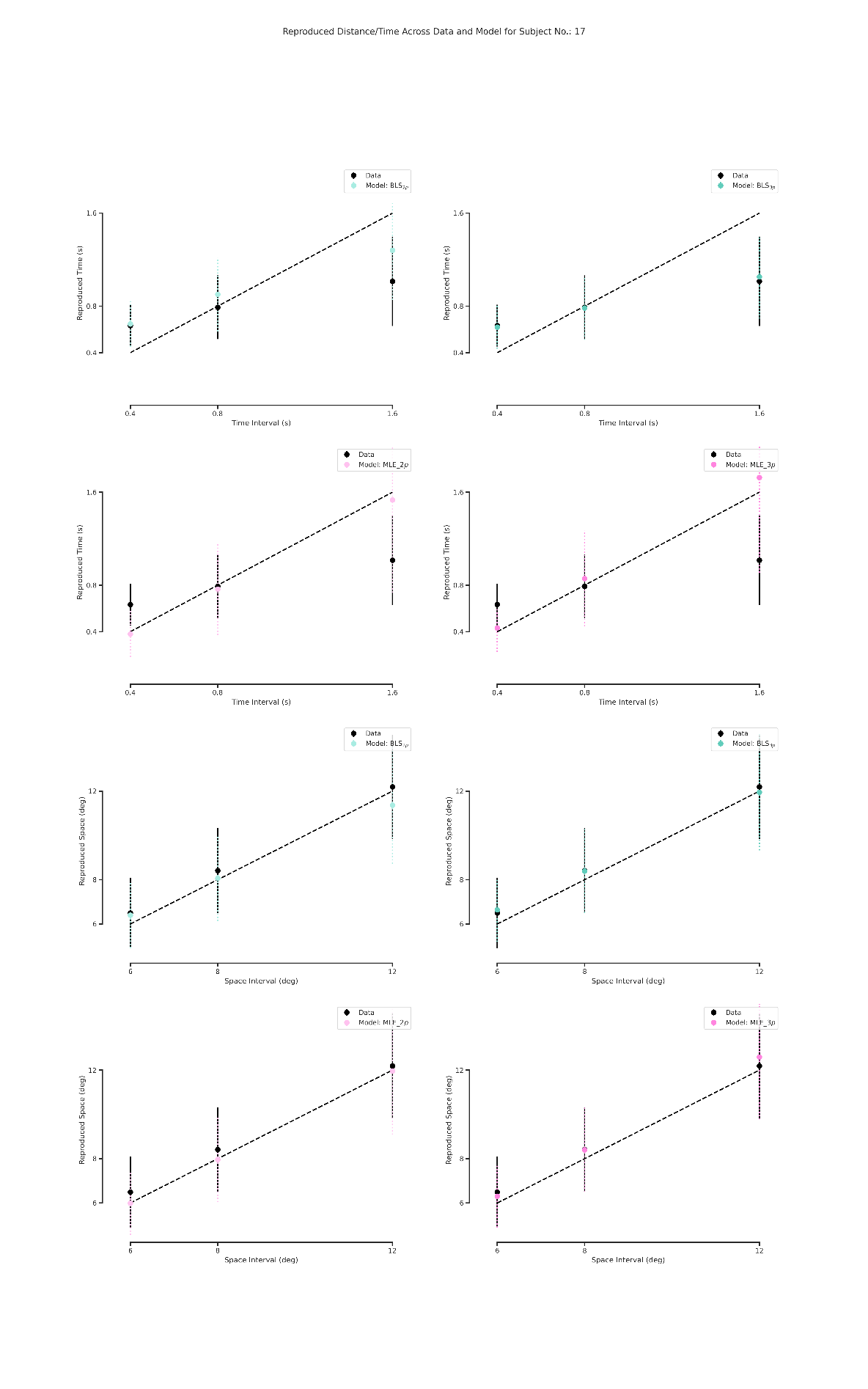

**Figure S17.** Subject and observer model behavior in time reproduction (Up) and in distance reproduction (Down).

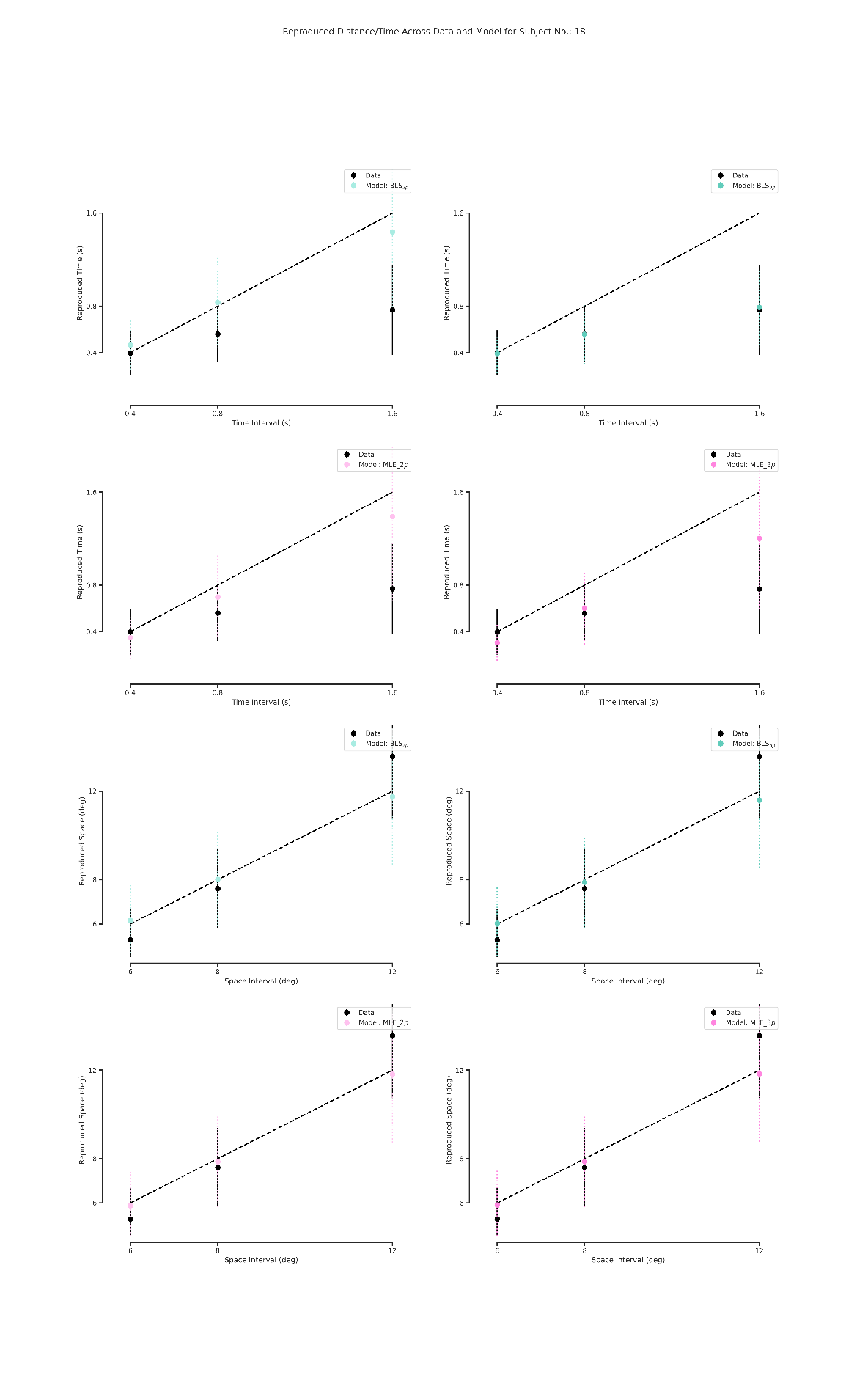

**Figure S18.** Subject and observer model behavior in time reproduction (Up) and in distance reproduction (Down).
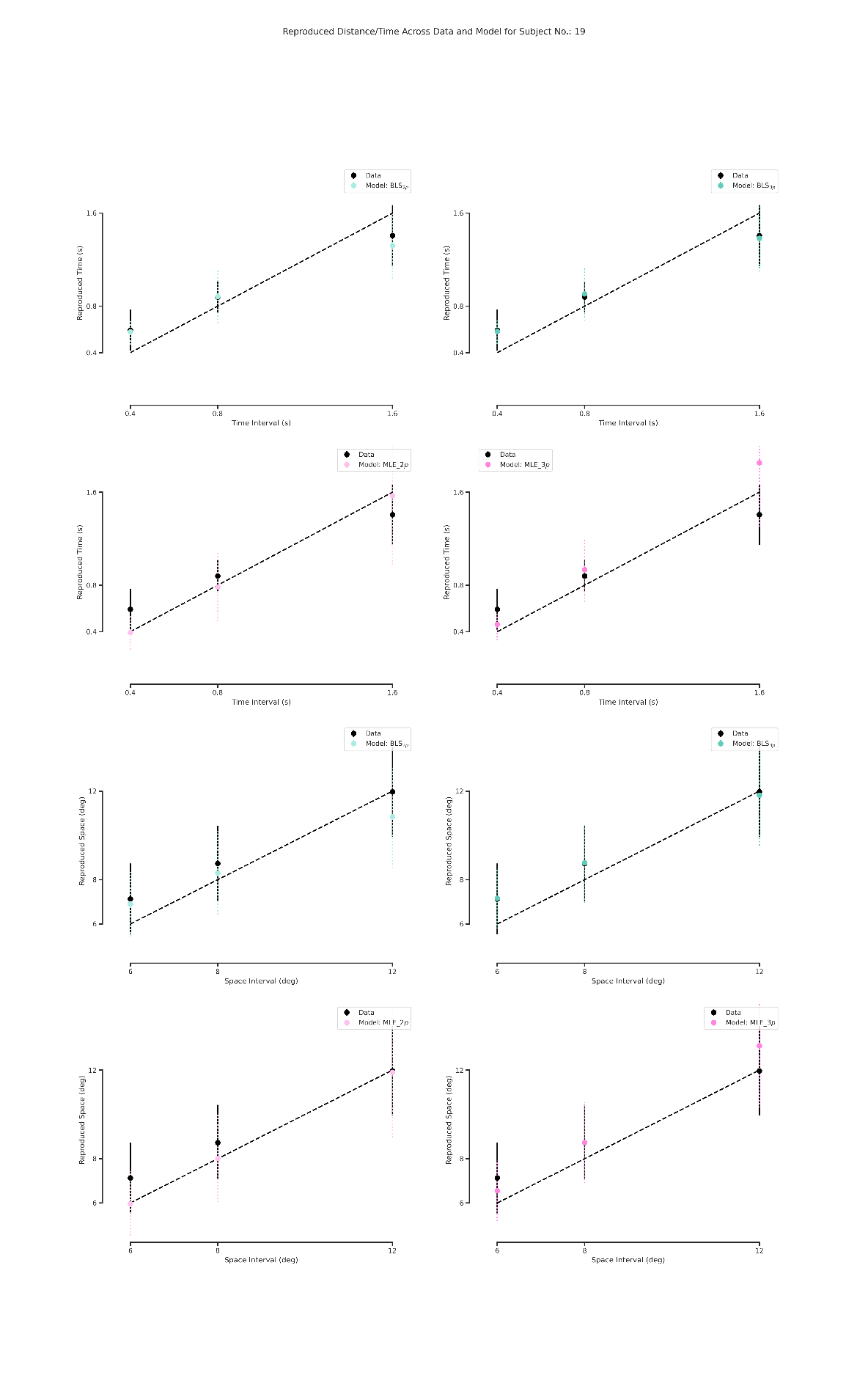

**Figure S19.** Subject and observer model behavior in time reproduction (Up) and in distance reproduction (Down).

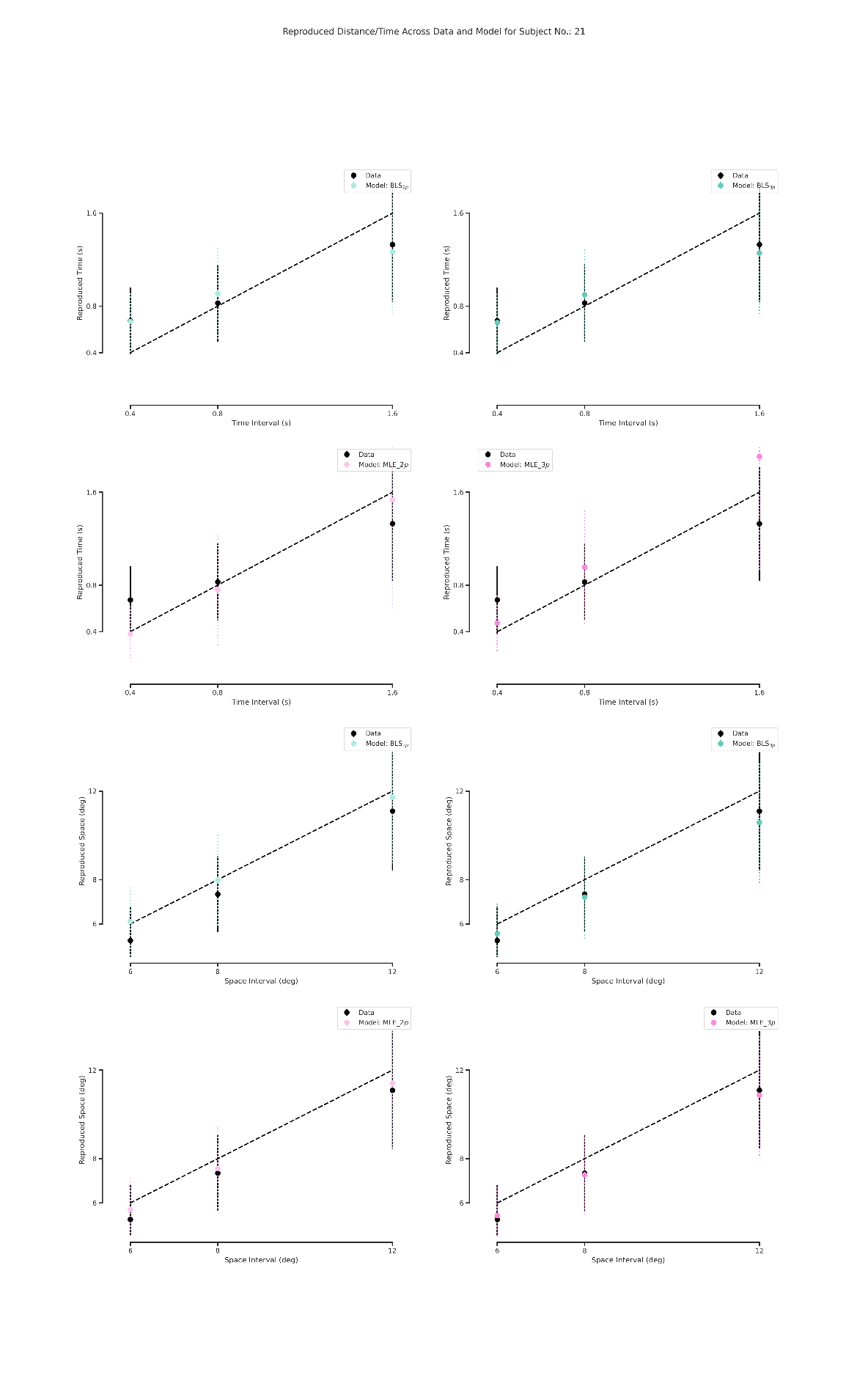

**Figure S20.** Subject and observer model behavior in time reproduction (Up) and in distance reproduction (Down).
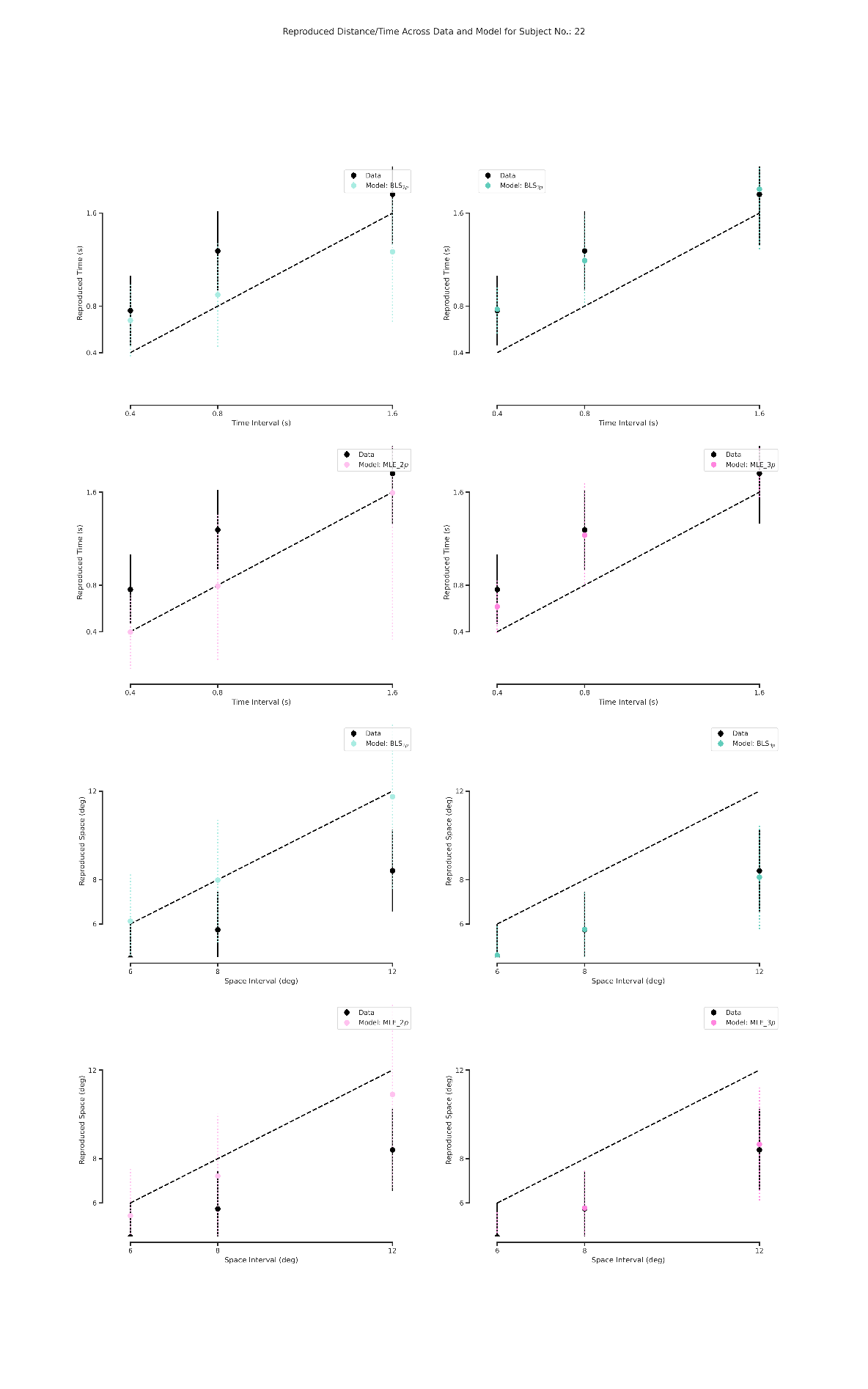
